# Automated model discovery for human brain using Constitutive Artificial Neural Networks

**DOI:** 10.1101/2022.11.08.515656

**Authors:** Kevin Linka, Sarah St. Pierre, Ellen Kuhl

## Abstract

The brain is our softest and most vulnerable organ, and understanding its physics is a challenging but significant task. Massive efforts have been dedicated at testing the human brain, and various competing models have emerged to characterize its response to mechanical loading. However, selecting the best constitutive model remains a heuristic process that strongly depends on user experience and personal preference. Here we challenge the conventional wisdom to first select a constitutive model and then fit its parameters to experimental data. Instead, we propose a new strategy that simultaneously discovers both model and parameters that best describe the data. Towards this goal, we integrate more than a century of knowledge in thermodynamics and state-of-the-art machine learning to build a family of Constitutive Artificial Neural Networks that enable automated model discovery for human brain tissue. Our overall design paradigm is to reverse engineer a Constitutive Artificial Neural Network from a set of functional building blocks that are, by design, a generalization of widely used and commonly accepted constitutive models, including the neo Hooke, Blatz Ko, Mooney Rivlin, Demiray, Gent, and Holzapfel models. By constraining the input, output, activation functions, and architecture, our network a priori satisfies thermodynamic consistency, material objectivity, material symmetry, physical constrains, and polyconvexity. We demonstrate that our network autonomously discovers both model and parameters that best characterize the behavior of human gray and white matter under tension, compression, and shear. Importantly, our network weights translate naturally into physically meaningful material parameters, e.g., shear moduli of 1.82kPa, 0.88kPa, 0.94kPa, and 0.54kPa for the cortex, basal ganglia, corona radiata, and corpus callosum. Our results suggest that Constitutive Artificial Neural Networks have the potential to induce a paradigm shift in soft tissue modeling, from user-defined model selection to automated model discovery. Our source code, data, and examples are available at https://github.com/LivingMatterLab/CANN.

## 1 Motivation

Traumatic brain injury is a major cause of death and disability worldwide [21], with a global annual incidence of 69 million [18]. In the United States alone, 176 people die each day from traumatic brain injury, and every nine seconds, someone sustains a new injury to the brain. Fortunately, not all concussions are life-threatening; yet, more than 5 million Americans are living with brain-injury-related disabilities and need long-term assistance in their everyday life [14]. Without a doubt, understanding the mechanics of the brain during traumatic brain injury is a challenging but significant task [25]. Throughout the past decade, scientists across the world have made significant strides in testing, modeling, and simulating the human brain [5, 10–12, 31, 49, 51, 61, 62]. However, because of its ultrasoft nature, the results vary greatly, both qualitatively and quantitatively [13]. This has resulted in a wide selection of competing constitutive models for gray and white matter tissue, without any real guidance of which model to choose [11, 41, 42, 45]. Throughout this manuscript, we ask how we can select the best constitutive model for the human brain, whether the current existing models are really the best, and if not, how we can systematically search and find a better model.

In machine learning, the process of finding relationships in complex data is known as *automated model discovery* [4, 20, 39]. The preface automated implies that model discovery can be performed entirely without human interaction [9, 54]. Neural networks have emerged as a powerful strategy to discover constitutive models from large data, even in the complete absence of knowledge about the underlying physics [30]. However, classical neural networks ignore more than a century of research in constitutive modeling [1]: They violate thermodynamic constraints [40], neglect generally accepted physical principles [3], and fail to predict the behavior outside the training regime [39]. In essence, neural networks perform excellently at fitting a complex function to big data, but they are not interpretable; they teach us nothing about the underlying physics [50]. So, really, what we are looking for is a strategy to autonomously *discover a physically motivated model*. Two successful but fundamentally different strategies have emerged to integrate physics into neural network models: *Physics-Informed Neural Networks* that add physics-based equations as additional terms to the loss function [33] and *Constitutive Artificial Neural Networks* that explicitly modify the network input, output, and architecture to hardwire physics-based constraints into the network design [36]. The first type of networks is more general and works well for ordinary [37] or partial [50] differential equations, whereas the second type is specifically tailored towards constitutive equations [38]. Constitutive Artificial Neural Networks, with strain invariants as input and free energy functions as output, were first proposed for rubber-like materials almost two decades ago [56], and have recently regained attention in the constitutive modeling community [3, 24, 34, 63]. They are now also increasingly recognized in the soft tissue biomechanics community with applications to skin [57], blood clots [32], arteries [29, 38], and myocardial tissue [32]. A common feature of all these neural networks is to use multiple hidden layers, generic activation functions, and several hundreds, if not thousands of unknowns. To no surprise, they perform well at interpolating non-linear stress-stretch relations from tension, compression, or shear experiments. However, one critical limitation remains: the lack of an intuitive interpretation of the model and its parameters [34].

Here, instead of using a generic neural network architecture, we *reverse-engineer* a new family of Constitutive Artificial Neural Networks from *constitutive building blocks* that are, by design, a generalization of widely used and commonly accepted constitutive models, including the neo Hooke [60], Blatz Ko [8], Mooney Rivlin [44, 52], Demiray [17], Gent [22], and Holzapfel [27] models. As such, their network weights naturally translate into material parameters with standard physical units and a clear physical interpretation [39]. We train our network with tension, compression, and shear tests from the human cortex, basal ganglia, corona radiata, and corpus callosum [11–13] and demonstrate that it can simultaneously *discover both model and parameters* that best describe the data. Beyond automated model discovery, we show that we can also use our network for the *parameter identification* of existing constitutive models. By systematically comparing the goodness of fit of the different models, trained with the different experiments, we not only discover the model that best describes the experiments, but we also *discover the experiment* that best informs the models. Designing informative experiments is particularly significant for human brain tissue, for which fresh samples are rare, challenging to preserve, and difficult to mount and test [13].

## 2 Methods

### 2.1 Kinematics

To characterize the deformation of the sample, we introduce the deformation map ***φ*** that maps material particles ***X*** from the undeformed configuration to particles, ***x*** = ***φ***(***X***), in the deformed configuration [2]. We describe relative deformations within the sample using the deformation gradient ***F***, the gradient of the deformation map ***φ*** with respect to the undeformed coordinates ***X***, and its Jacobian *J*,

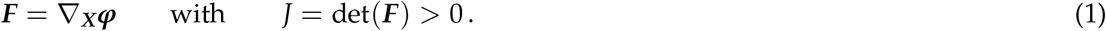

Multiplying ***F*** with its transpose ***F***^t^ introduces the symmetric right Cauchy Green deformation tensor ***C***,

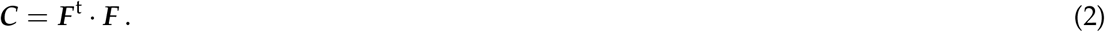

In the undeformed state, both tensors are identical to the unit tensor, ***F*** = ***I*** and ***C*** = ***I***, and the Jacobian is one, *J* = 1. A Jacobian smaller than one, 0 *< J <* 1, denotes compression and a Jacobian larger than one, 1 *< J*, denotes extension.

#### Isotropy

To characterize an *isotropic* material, we introduce the three principal invariants *I*_1_, *I*_2_, *I*_3_ and their derivatives *∂*_***F***_ *I*_1_, *∂*_***F***_ *I*_2_, *∂*_***F***_ *I*_3_,

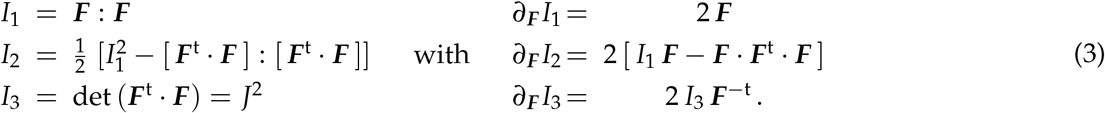

In the undeformed state, ***F*** = ***I***, the three invariants are equal to three and one, *I*_1_ = 3, *I*_2_ = 3, and *I*_3_ = 1.

#### Perfect incompressibility

For *isotropic, perfectly incompressible* materials, the third invariant always remains identical to one, *I*_3_ = *J*^2^ = 1. This reduces the set of invariants to two, *I*_1_ and *I*_2_.

### 2.2 Constitutive equations

In solid mechanics, constitutive equations are tensor-valued tensor functions that define the relation between a stress, for example the Piola or nominal stress, ***P*** = lim_d*→* ***A*0**_ (d ***f*** /d***A***), the force d ***f*** per undeformed area d***A***, and a deformation measure, for example the deformation gradient ***F*** [28, 58],

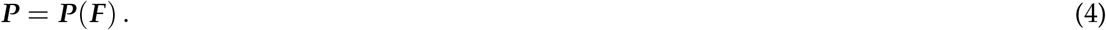

At this point, we could use an arbitrary neural network to learn the functional relation between ***P*** and ***F*** and many neural networks in the literature do exactly that [23, 40, 55]. However, the functions ***P***(***F***) that we learn through this approach generally violate widely-accepted thermodynamical constraints and their parameters have no physical meaning [24]. For moderate amounts of data, standard neural networks are also associated with a high risk of overfitting [34]. Our objective is therefore to build a Constitutive Artificial Neural Network that a priori satisfies thermodynamic constraints and introduces parameters with a clear physical interpretation, while, at the same time, limiting the space of admissible functions to prevent overfitting when available data are sparse.

#### Thermodynamic consistency

First, we ensure *thermodynamical consistency* and guarantee that the Piola stress ***P*** inherently satisfies the second law of thermodynamics, the dissipation inequality [48], 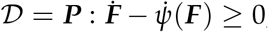, where 𝒟 is the dissipation and *ψ* is the Helmholtz free energy with 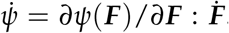. For *hyperelastic* or Green-elastic materials with 𝒟 ≐ 0, the entropy inequality defines the Piola stress [59],

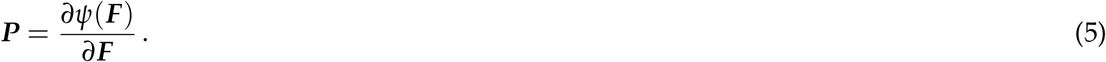

This implies that, rather than approximating the nine stress components ***P***(***F***) as nine generic functions of the nine components of the deformation gradient ***F***, the network approximates the free energy function *ψ*(***F***) from which we derive the stress ***P*** in a post-processing step. Satisfying thermodynamic consistency according to equation (5) directly affects the *output* of the neural network.

#### Material objectivity and frame indifference

Second, we constrain the choice of the free energy function *ψ* to satisfy *material objectivity* or *frame indifference* and ensure that the constitutive laws do not depend on the external frame of reference [46]. This implies that the arguments of the free energy function are independent of rotations and must be functions of the right Cauchy Green deformation tensor ***C*** [58],

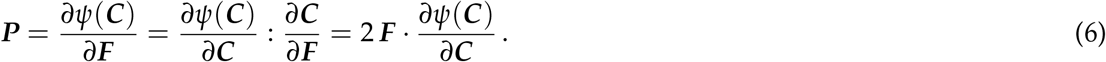

This implies that, rather than using the nine independent components of the deformation gradient ***F*** as input, we constrain the input to the six independent components of the symmetric right Cauchy Green deformation tensor, ***C*** = ***F***^t^ · ***F***. Satisfying material objectivity according to equation directly affects the *input* of the neural network.

#### Material symmetry and isotropy

Third, we can further constrain the choice of the free energy function *ψ* to include *material symmetry* and assume that the material response remains unchanged under transformations of the reference configuration. Here we consider the special case of *isotropy*, for which the free energy function is a function of the strain *invariants, ψ*(*I*_1_, *I*_2_, *I*_3_), and the Piola stress takes the following explicit representation,

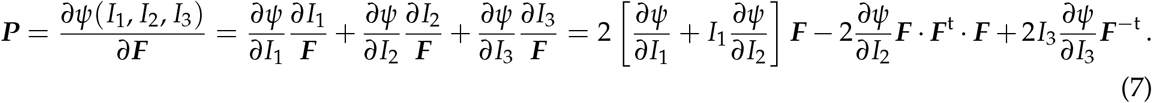

This implies that, rather than using the six independent components of the symmetric right Cauchy Green deformation tensor ***C*** as input, we constrain the input to our set of three invariants *I*_1_, *I*_2_, *I*_3_. Considering materials with known symmetry classes according to equations (7) directly affects, and ideally reduces, the *input* of the neural network.

#### Perfect incompressibility

Fourth, we can further constrain the choice of the free energy function *ψ* for the special case of *perfect incompressibility* for which the Jacobian remains constant and equal to one, *I*_3_ = *J*^2^ = 1. The condition of perfect incompressibility implies that equation (7) simplifies to an expression in terms of only the first two invariants *I*_1_ and *I*_2_, supplemented by a term of the hydrostatic pressure, 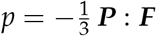, that we determine from the boundary conditions,

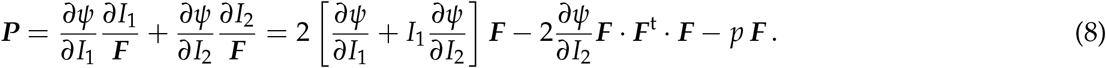

This implies that, rather than using the set of three invariants, *I*_1_, *I*_2_, *I*_3_, as input, we reduce the input to a set of only two invariants, *I*_1_ and *I*_2_. Considering materials with perfect incompressibility according to equation (8) reduces the *input* of the neural network.

#### Physically reasonable constitutive restrictions

Fifth, we can further constrain the functional form of the free energy *ψ* by including additional constitutive restrictions that are both physically reasonable and mathematically convenient [2]:

i. The free energy *ψ* is *non-negative* for all deformation states ***F***,

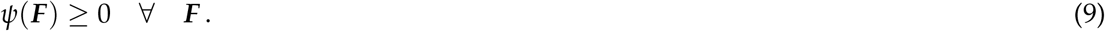
ii. The free energy *ψ* and the stress ***P*** are *zero* in the reference configuration, ***F*** = ***I***,

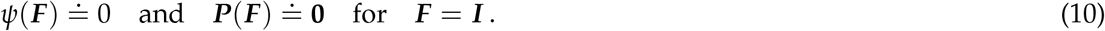
iii. The free energy *ψ* is *infinite* for infinite compression, *J →* 0, and infinite expansion, *J →* ∞,

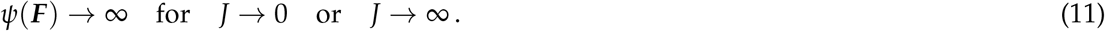

To facilitate a stress-free reference configuration according to equation (10), instead of using the invariants *I*_1_, *I*_2_, *I*_3_ themselves as input, we use their deviation from the energy- and stress-free reference state, [*I*_1_ − 3], [*I*_2_ − 3], [*I*_3_ − 1], as input. In addition, from all possible activation functions, we select activation functions that a priori comply with conditions (i), (ii), and (iii). Satisfying physical considerations according to equations (9), (10), and (11) directly affects the *activation functions* of the neural network.

#### Polyconvexity

Sixth, to guide the selection of the functional forms for the free energy function *ψ*, and ultimately the selection of appropriate activation functions, we consider *polyconvexity* requirements [6]. From the general representation theorem we know that in its most generic form, the free energy of an isotropic material can be expressed as an infinite series of products of powers of the invariants [53], 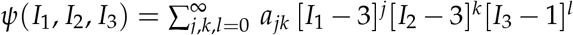, where *a*_*jkl*_ are material constants. Importantly, mixed products of convex functions are generally not convex, and it is easier to show that the sum of specific convex subfunction usually is [26]. This motivates a special subclass of free energy functions in which the free energy is the sum of three individual polyconvex subfunctions *ψ*_1_, *ψ*_2_, *ψ*_3_, such that *ψ*(***F***) = *ψ*_1_(*I*_1_) + *ψ*_2_(*I*_2_) + *ψ*_3_(*I*_3_), is polyconvex by design and the stresses take the following form,

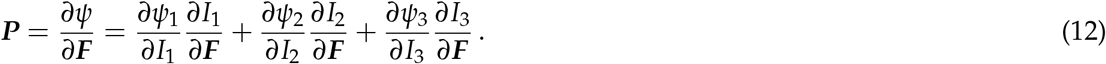

This implies that we can either select polyconvex activation functions from a set of algorithmically predefined activation functions [39], or custom-design our own activations functions from known polyconvex subfunctions *ψ*_1_, *ψ*_2_, *ψ*_3_ [3]. Here we select first and second powers of the invariants for the first hidden layer and linear, exponential, and logarithmic functions of these powers for the second hidden layer, all with *non-negative coefficients*. In addition, we abandon the fully-connected network architecture, in which mixed products of the invariants *I*_1_, *I*_2_, *I*_3_ emerge naturally. Instead, we decouple the inputs *I*_1_, *I*_2_, *I*_3_ and only combine them additively in the free energy function, *ψ* = *ψ*_1_ + *ψ*_2_ + *ψ*_3_. Satisfying polyconvexity, for example according to equation (12), can imply enforcing *non-negative network weights* [3], and directly affects the *architecture* of the neural network [34].

### 2.3 Constitutive Artificial Neural Networks

Motivated by these considerations, we build a family of Constitutive Artificial Neural Networks that satisfy the conditions of thermodynamic consistency, material objectivity, material symmetry, incompressibility, constitutive restrictions, and polyconvexity by design. This guides our selection of network *input, output, architecture*, and *activation functions* to a priori satisfy the fundamental laws of physics. Special members of this family represent well-known constitutive models, including the neo Hooke [60], Blatz Ko [8], Mooney Rivlin [44, 52], Demiray [17], Gent [22], and Holzapfel [27] models, for which the network weights gain a clear physical interpretation.

#### Constitutive Artificial Neural Network input and output

To ensure thermodynamical consistency, rather than directly approximating the stress ***P*** as a function of the deformation gradient ***F***, the Constitutive Artificial Neural Network approximates the scalar-valued free energy function *ψ* as a function of the scalar-valued invariants *I*_1_, *I*_2_, *I*_3_. The Piola stress ***P*** then follows naturally from the second law of thermodynamics as the derivative of the free energy *ψ* with respect to the deformation gradient ***F*** according to equation (7). Figure 1 illustrates a Constitutive Artificial Neural Network with the invariants *I*_1_, *I*_2_, *I*_3_ as input and the the free energy *ψ* as output.

**Figure 1:**
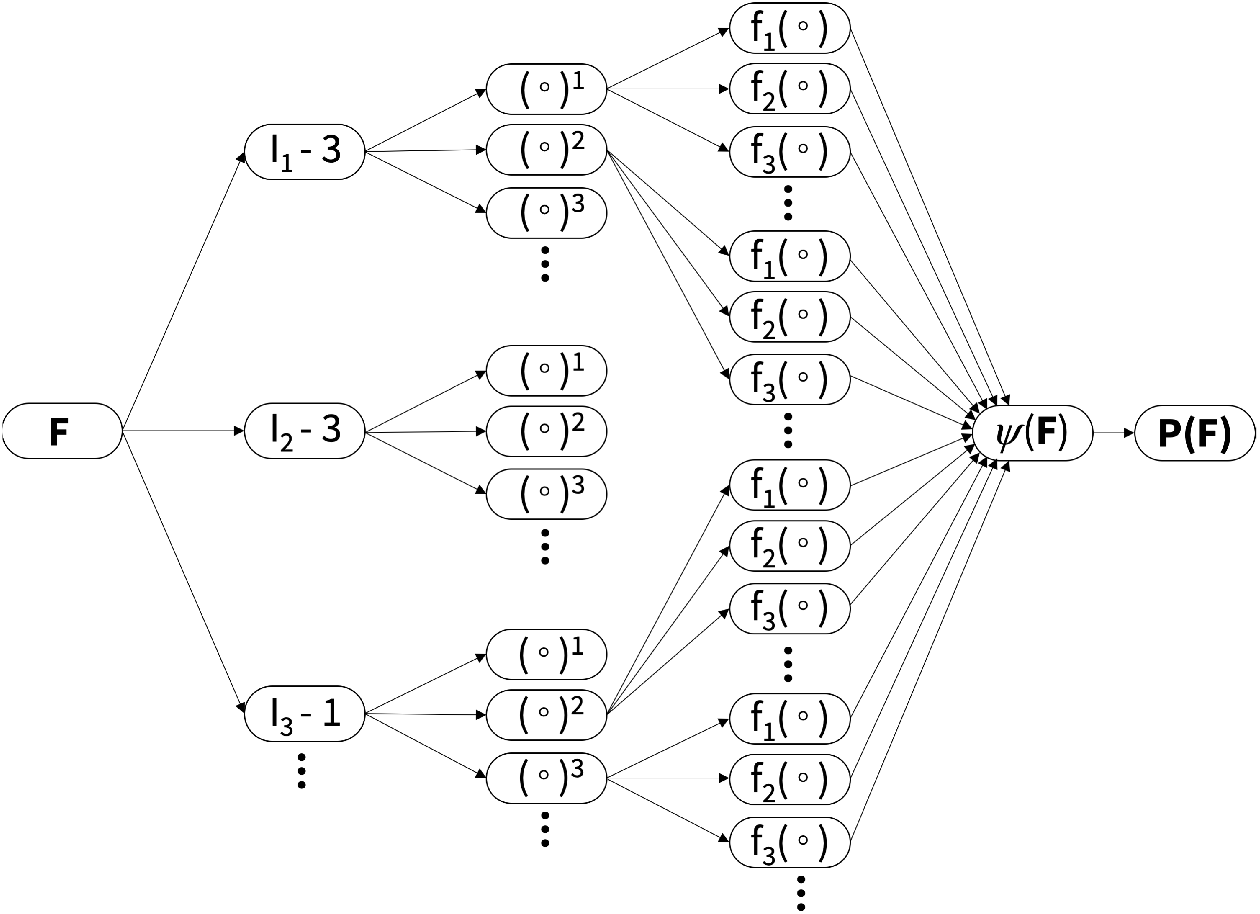
Constitutive Artificial Neural Network. Family of a feed forward Constitutive Artificial Neural Networks with two hidden layers to approximate the single scalar-valued free energy function *ψ*(*I*_1_, *I*_2_, *I*_3_) as a function of the scalarvalued invariants *I*_1_, *I*_2_, *I*_3_ of the deformation gradient ***F***. The first layer generates powers (∘), (∘)_2_, (∘)_3_ of the network input and the second layer applies thermodynamically admissible activation functions *f* (∘) to these powers. Constitutive Artificial Neural Networks are typically not fully connected by design to a priori satisfy the condition of polyconvexity.

#### Constitutive Artificial Neural Network architecture

To model a hyperelastic *history-independent* material, we select a feed forward architecture in which information only moves in one direction, from the input nodes, without any cycles or loops, to the output nodes. To ensure polyconvexity, we choose a selectively connected architecture according to equation (12), such that the free energy function does not contain mixed terms in the invariants. Figure 1 illustrates one possible network architecture that a priori decouples the individual invariants. Its free energy function,

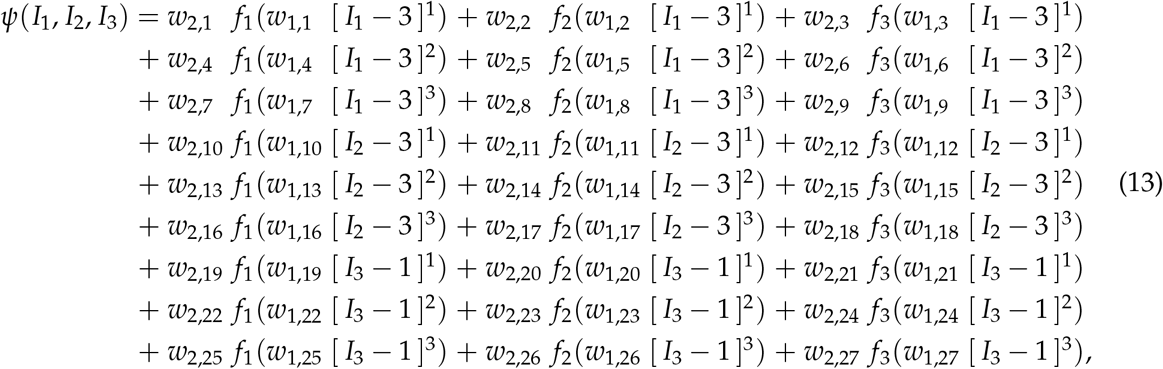

introduces 3 *×* 3 *×* 3 + 3 *×* 3 *×* 3 = 54 weights. The first set of 27 weights, *w*_1,1..27_, weighs the powers of the invariants and the second set of 27 weights, *w*_2,1..27_, weighs the contributions of the functions *f*_1_, *f*_2_, *f*_3_.

#### Activation functions

To ensure that our network satisfies basic physically reasonable constitutive restrictions, rather than selecting from a set of pre-defined activation functions such as the binary step, soft step, hyperbolic tangent, inverse tangent, or soft plus functions, we custom-design our own activation functions to *reverse-engineer* a free energy function that captures popular forms of constitutive terms. Specifically, we select linear and quadratic powers of the first and second invariants for the first layer of the network, and linear, exponential, or logarithmic functions for the second layer.

Figure 2 illustrates the six activation functions *f* (*x*) along with their derivatives *f* (*x*) that we use throughout the remainder of this work. Notably, in contrast to the activation functions for classical neural networks, all six functions are not only *monotonic, f* (*x* + *ε*) ≥ *f* (*x*) for *ε* ≥ 0, such that increasing deformations result in increasing stresses, but also *continuous* at the origin, *f* (−0) = *f* (+0), *continuously differentiable* and *smooth* at the origin, *f* (−0) = *f* (+0), and zero at the origin, *f* (0) = 0, to ensure an energy- and stress-free reference configuration according to equation (10), and *unbounded, f* (−∞) *→* ∞ and *f* (+∞)*→* ∞, to ensure an infinite energy and stress for extreme deformations according to equation (11).

**Figure 2:**
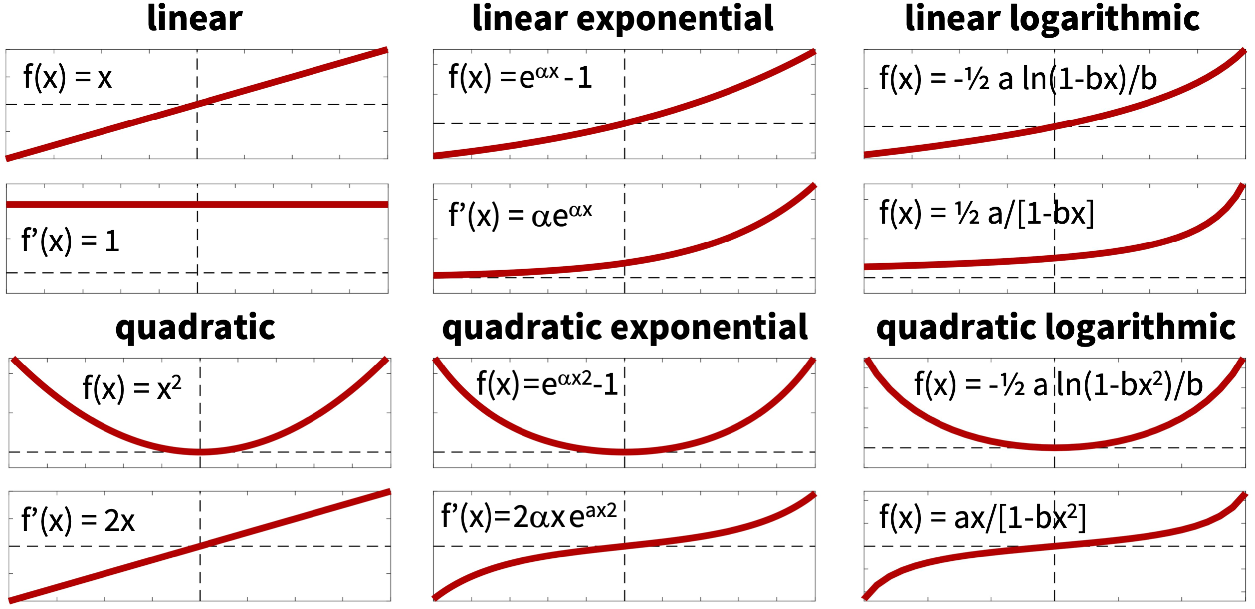
Activation functions for Constitutive Artificial Neural Networks. We use custom-design activation functions *f* (*x*) along with their derivatives *f* (*x*) that include linear and quadratic mappings, left, linear and quadratic exponentials, middle, and linear and quadratic logarithmic functions, right, to reverse engineer a free energy function that captures popular functional forms of constitutive terms.

Figure 3 illustrates our isotropic, perfectly incompressible Constitutive Artificial Neural Network with two hidden layers and four and twelve nodes. The first layer generates powers (*∘*)^1^ and (∘)^2^ of the network inputs, [*I*_1_ − 3] and [*I*_2_ − 3], and the second layer applies the identity, (∘), the exponential function, (exp((*∘*))− 1), and the natural logarithm, (−ln(1 − (*∘*))), to these powers. The set of equations for this networks takes the following explicit form,

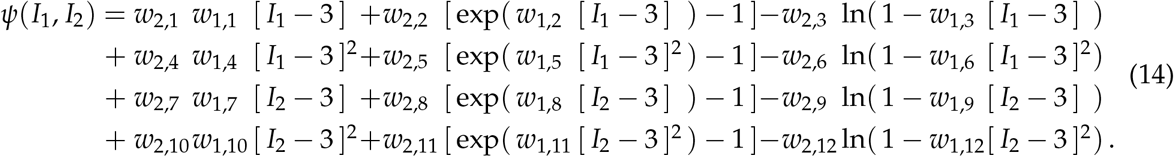

For this particular format, one of the first two weights of each row becomes redundant, and we can reduce the set of network parameters from 24 to 20, **w** = [(*w*_1,1_ *w*_2,1_), *w*_1,2_, *w*_2,2_, *w*_1,3_, *w*_2,3_, (*w*_1,4_ *w*_2,4_), *w*_1,5_, *w*_2,5_, *w*_1,6_, *w*_2,6_, (*w*_1,7_ *w*_2,7_), *w*_1,8_, *w*_2,8_, *w*_1,9_, *w*_2,9_, (*w*_1,10_ *w*_2,10_), *w*_1,11_, *w*_2,11_, *w*_1,12_, *w*_2,12_]. Using the second law of thermodynamics, we can derive an explicit expression for the Piola stress from equation (5), ***P*** = *∂ψ*/*∂****F***, or, more specifically, for the case of perfect incompressibility from equation (8), ***P*** = *∂ψ*/*∂I*_1_ *· ∂I*_1_/*∂****F*** + *∂ψ*/*∂I*_2_ *· ∂I*_2_/*∂****F***,

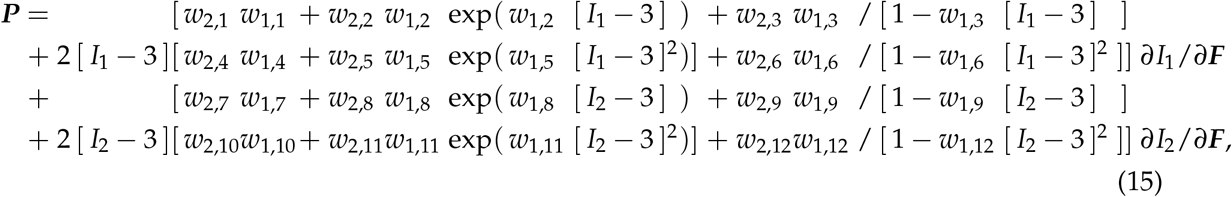

corrected by the pressure term, 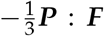. The stress definition (15) suggests that our model is a *generalization* of many popular constitutive models for incompressible hyperelastic materials. It seems natural to ask whether and how its network parameters *w*_1..2,1..12_ relate to the parameters of these models.

**Figure 3:**
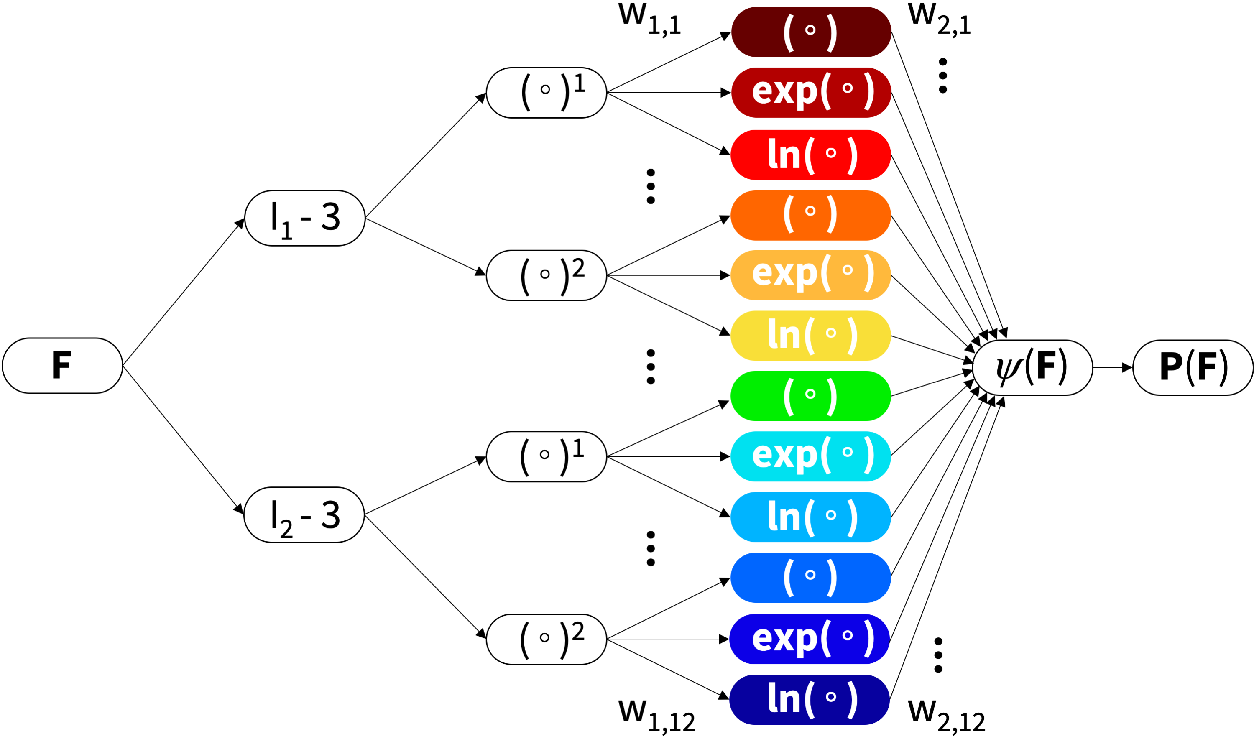
Constitutive Artificial Neural Network. Isotropic, perfectly incompressible Constitutive Artificial Neural Network with with two hidden layers to approximate the single scalar-valued free energy function *ψ*(*I*_1_, *I*_2_) as a function of the first and second invariants of the deformation gradient ***F*** using twelve terms. The first layer generates powers (*∘*)^1^ and (*∘*)^2^ of the network inputs, [*I*_1_ − 3] and [*I*_2_ − 3] and the second layer applies the identity (∘), the exponential function, (exp((*∘*))− 1), and the natural logarithm, (−ln(1 − (∘))), to these powers. The networks is selectively connected by design to a priori satisfy the condition of polyconvexity.

#### Special types of constitutive equations

To demonstrate that the family of Constitutive Artificial Neural Networks in Figure 1 and its specific example in Figure 3 are *generalizations* of popular constitutive models, we consider six widely used models and systematically compare their material parameters to our network weights *w*_1..2,1..12_:

The *neo Hooke model* [60], the simplest of all models, has a free energy function that is a constant function of only the first invariant, [*I*_1_ − 3], scaled by the shear modulus *μ*,

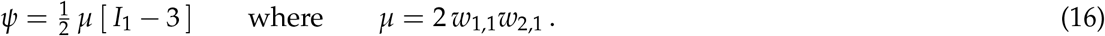

The *Blatz Ko model* [8], has a free energy function that depends only the second and third invariants, [*I*_2_ − 3] and [*I*_3_ − 1], scaled by the shear modulus *μ* as 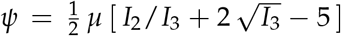. For perfectly incompressible materials, *I*_3_ = 1, it simplifies to the following form,

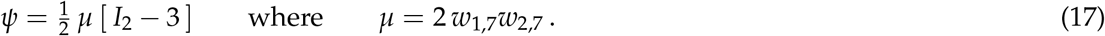

The *Mooney Rivlin model* [44, 52] is a combination of both and accounts for the first and second invariants, [*I*_1_ − 3] and [*I*_2_ − 3], scaled by the moduli *μ*_1_ and *μ*_2_ that sum up to the overall shear modulus, *μ* = *μ*_1_ + *μ*_2_,

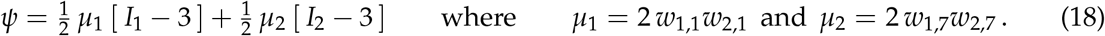

The *Demiray model* [17] or *Delfino model* [16] uses linear exponentials of the first invariant, [*I*_1_ − 3], in terms of two parameters *a* and *b*,

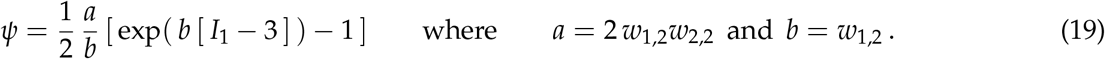

The *Gent model* [22] uses linear logarithms of the first invariant, [*I*_1_ − 3], in terms of two parameters *α* and *β*,

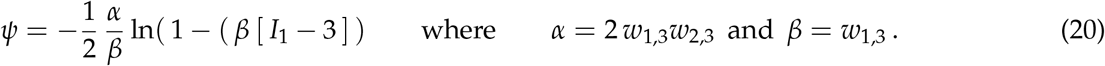

The *Holzapfel model* [27] uses quadratic exponentials, typically of the fourth invariant, which we adapt here for the the first invariant, [*I*_1_ − 1], in terms of two parameters *a* and *b*,

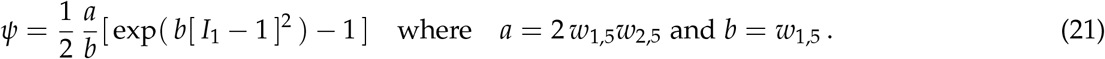

These simple examples demonstrate that we can recover popular constitutive functions for which the network weights gain a well-defined physical meaning.

#### Loss function

The objective of our Constitutive Artificial Neural Network is to learn the network parameters ***θ*** = {*w*_*ij*_}, the network weights, by minimizing a loss function *L* that penalizes the error between model and data. Similar to classical neural networks, we characterize this error as the mean squared error, the *L*_2_-norm of the difference between model ***P***(***F***_*i*_) and data 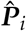, divided by the number of training points *n*_trn_,

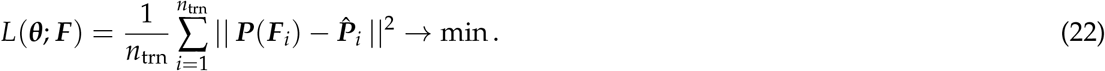

To reduce potential overfitting, we also study the effects of Lasso or L1 regularization and L2 regularization,

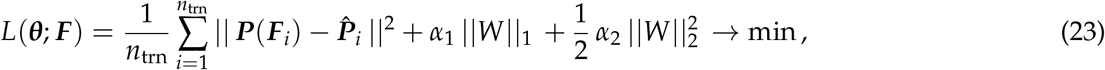

by enhancing the loss function by the weighted L1 norm, ||*W*||_1_ = ∈_*i*_ ∈_*j*_ | *w*_*ij*_ |, or the weighted Euclidian or L2 norm, 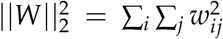, where *α*_1_ and *α*_2_ are the weighting coefficients. We train the network by minimizing the loss function (22) or (23) and learn the network parameters ***θ*** = *{w*_*ij*_*}* using the ADAM optimizer, a robust adaptive algorithm for gradient-based first-order optimization, and constrain the weights to always remain non-negative, *w*_*ij*_ ≥ 0.

### 2.4 Data

To illustrate the automated model discovery with our Constitutive Artificial Neural Network, we perform a systematic study using widely-used benchmark data for human brain tissue [11–13]. Specifically, we train our two-layer Constitutive Artificial Neural Network for isotropic, perfectly incompressible materials from Figure 3, discover a material model and its parameters, and compare the model and parameters against six traditional constitutive models for soft biological tissues [8,17,22,27,44,52,60]. We consider two training scenarios, *single-mode training* and *multi-mode training*, for the homogeneous deformation modes of uniaxial tension, uniaxial compression, and simple shear.

#### Tension and compression

For the case of uniaxial tension and compression, we stretch the specimen in one direction, *F*_11_ = *λ*_1_ = *λ*. For an *isotropic, perfectly incompressible* material with 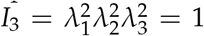, the stretches orthogonal to the loading direction are identical and equal to the square root of the stretch, *F*_22_ = *λ*_2_ = *λ*^−1/2^ and *F*_33_ = *λ*_3_ = *λ*^−1/2^. From the resulting deformation gradient, ***F*** = diag *λ, λ*^−1/2^, *λ*^−1/2^, we calculate the first and second invariants and their derivatives,

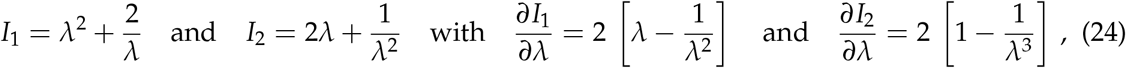

to evaluate the nominal uniaxial stress *P*_11_ using the general stress-stretch relationship for perfectly incompressible materials, *P*_*ii*_ = [*∂ψ*/*∂I*_1_] [*∂I*_1_/*∂λ*_*i*_] + [*∂ψ*/*∂I*_2_] [*∂I*_2_/*∂λ*_*i*_] − [1/*λ*_*i*_] *p*, for *i* = 1, 2, 3. Here, *p* denotes the hydrostatic pressure that we determine from the zero stress condition in the transverse directions, *P*_22_ = 0 and *P*_33_ = 0, as *p* = [2/*λ*] *∂ψ*/*∂I*_1_ + [2*λ* + 2/*λ*^2^] *∂ψ*/*∂I*_2_. This results in the following explicit *uniaxial stress-stretch relation for perfectly incompressible, isotropic* materials,

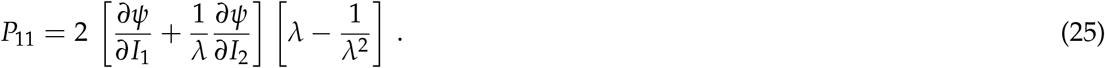

#### Shear

For the case of simple shear, we shear the specimen in one direction, *F*_12_ = *γ*. For an *isotropic, perfectly incompressible* material with *F*_11_ = *F*_22_ = *F*_33_ = 1, we calculate the first and second invariants and their derivatives,

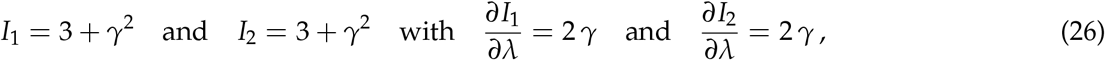

to evalute the nominal shear stress *P*_12_ using the general stress-stretch relationship for perfectly incompressible materials. This results in the following explicit *shear stress-strain relation for perfectly incompressible, isotropic* materials,

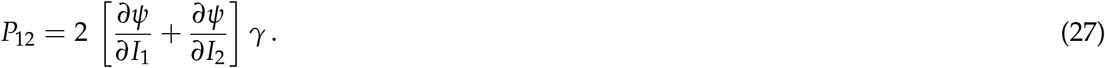

#### Testing and training data

Tables 1 and 2 summarize our benchmark data of gray matter tissue from the cortex and basal ganglia and white matter tissue from the corona radiata and corpus callosum tested in tension, compression, and shear [11–13]. We report each data set as 17 pairs of stretches and nominal uniaxial stresses, {*λ, P*_11_} or shear strains and nominal shear stresses, {*γ, P*_12_}, where the stretches and shear strains range from 0.9 ≤ *λ* ≤ 1.1 and 0.0 ≤ *γ* ≤ 0.2, and the stresses are the means of the loading and unloading curves of *n* samples. Throughout the remainder of this study, we perform *single-mode training* with one mode used as training data and the remaining two modes as test data, and *multi-mode training* with all three modes used as training data.

**Table 1:** Gray matter data. Cortex and basal ganglia tested in tension, compression, and shear; stresses are reported as means from the loading and unloading curves of *n* samples [11].

| cortex tension<br>$n = 15$ | | cortex compression<br>$n = 17$ | | cortex shear<br>$n = 35$ | | basal ganglia tension<br>$n = 15$ | | basal ganglia compression<br>$n = 15$ | | basal ganglia shear<br>$n = 29$ | |
| --- | --- | --- | --- | --- | --- | --- | --- | --- | --- | --- | --- |
| $\lambda$<br>[-] | $P_{11}$<br>[kPa] | $\lambda$<br>[-] | $P_{11}$<br>[kPa] | $\gamma$<br>[-] | $P_{12}$<br>[kPa] | $\lambda$<br>[-] | $P_{11}$<br>[kPa] | $\lambda$<br>[-] | $P_{11}$<br>[kPa] | $\gamma$<br>[-] | $P_{12}$<br>[kPa] |
| 1.0000 | 0.0000 | 1.0000 | 0.0000 | 0.0000 | 0.0000 | 1.0000 | 0.0000 | 1.0000 | 0.0000 | 0.0000 | 0.0000 |
| 1.0063 | 0.0251 | 0.9938 | -0.0308 | 0.0125 | 0.0147 | 1.0063 | 0.0149 | 0.9938 | -0.0174 | 0.0125 | 0.0070 |
| 1.0125 | 0.0462 | 0.9875 | -0.0659 | 0.0250 | 0.0294 | 1.0125 | 0.0251 | 0.9875 | -0.0358 | 0.0250 | 0.0140 |
| 1.0188 | 0.0666 | 0.9812 | -0.1040 | 0.0375 | 0.0486 | 1.0188 | 0.0345 | 0.9812 | -0.0534 | 0.0375 | 0.0210 |
| 1.0250 | 0.0838 | 0.9750 | -0.1479 | 0.0500 | 0.0633 | 1.0250 | 0.0446 | 0.9750 | -0.0778 | 0.0500 | 0.0305 |
| 1.0312 | 0.1010 | 0.9688 | -0.1908 | 0.0625 | 0.0814 | 1.0312 | 0.0540 | 0.9688 | -0.1021 | 0.0625 | 0.0397 |
| 1.0375 | 0.1175 | 0.9625 | -0.2375 | 0.0750 | 0.0983 | 1.0375 | 0.0619 | 0.9625 | -0.1265 | 0.0750 | 0.0488 |
| 1.0437 | 0.1324 | 0.9563 | -0.2920 | 0.0875 | 0.1186 | 1.0437 | 0.0705 | 0.9563 | -0.1479 | 0.0875 | 0.0579 |
| 1.0500 | 0.1488 | 0.9500 | -0.3504 | 0.1000 | 0.1412 | 1.0500 | 0.0791 | 0.9500 | -0.1752 | 0.1000 | 0.0703 |
| 1.0562 | 0.1661 | 0.9437 | -0.4127 | 0.1125 | 0.1649 | 1.0562 | 0.0862 | 0.9437 | -0.2102 | 0.1125 | 0.0805 |
| 1.0625 | 0.1856 | 0.9375 | -0.4866 | 0.1250 | 0.1942 | 1.0625 | 0.0963 | 0.9375 | -0.2414 | 0.1250 | 0.0930 |
| 1.0688 | 0.2091 | 0.9313 | -0.5684 | 0.1375 | 0.2292 | 1.0688 | 0.1050 | 0.9313 | -0.2842 | 0.1375 | 0.1088 |
| 1.0750 | 0.2366 | 0.9250 | -0.6579 | 0.1500 | 0.2698 | 1.0750 | 0.1151 | 0.9250 | -0.3270 | 0.1500 | 0.1257 |
| 1.0813 | 0.2710 | 0.9187 | -0.7630 | 0.1625 | 0.3227 | 1.0813 | 0.1277 | 0.9187 | -0.3776 | 0.1625 | 0.1449 |
| 1.0875 | 0.3125 | 0.9125 | -0.8837 | 0.1750 | 0.3791 | 1.0875 | 0.1426 | 0.9125 | -0.4321 | 0.1750 | 0.1686 |
| 1.0938 | 0.3650 | 0.9062 | -1.0005 | 0.1875 | 0.4557 | 1.0938 | 0.1582 | 0.9062 | -0.4905 | 0.1875 | 0.1969 |
| 1.1000 | 0.4151 | 0.9000 | -1.1484 | 0.2000 | 0.5435 | 1.1000 | 0.1778 | 0.9000 | -0.5528 | 0.2000 | 0.2262 |

**Table 2:** White matter data. Corona radiata and corpus callosum tested in tension, compression, and shear; stresses are reported as means from the loading and unloading curves of *n* samples [11].

| corona radiata tension<br>$n = 18$ | | corona radiata compression<br>$n = 18$ | | corona radiata shear<br>$n = 36$ | | corpus callosum tension<br>$n = 19$ | | corpus callosum compression<br>$n = 20$ | | corpus callosum shear<br>$n = 39$ | |
| --- | --- | --- | --- | --- | --- | --- | --- | --- | --- | --- | --- |
| $\lambda$<br>[-] | $P_{11}$<br>[kPa] | $\lambda$<br>[-] | $P_{11}$<br>[kPa] | $\gamma$<br>[-] | $P_{12}$<br>[kPa] | $\lambda$<br>[-] | $P_{11}$<br>[kPa] | $\lambda$<br>[-] | $P_{11}$<br>[kPa] | $\gamma$<br>[-] | $P_{12}$<br>[kPa] |
| 1.0000 | 0.0000 | 1.0000 | 0.0000 | 0.0000 | 0.0000 | 1.0000 | 0.0000 | 1.0000 | 0.0000 | 0.0000 | 0.0000 |
| 1.0063 | 0.0157 | 0.9938 | -0.0193 | 0.0125 | 0.0079 | 1.0063 | 0.0078 | 0.9938 | -0.0096 | 0.0125 | 0.0036 |
| 1.0125 | 0.0235 | 0.9875 | -0.0387 | 0.0250 | 0.0159 | 1.0125 | 0.0149 | 0.9875 | -0.0164 | 0.0250 | 0.0072 |
| 1.0188 | 0.0345 | 0.9812 | -0.0543 | 0.0375 | 0.0238 | 1.0188 | 0.0196 | 0.9812 | -0.0300 | 0.0375 | 0.0109 |
| 1.0250 | 0.0423 | 0.9750 | -0.0800 | 0.0500 | 0.0318 | 1.0250 | 0.0251 | 0.9750 | -0.0427 | 0.0500 | 0.0170 |
| 1.0312 | 0.0509 | 0.9688 | -0.1040 | 0.0625 | 0.0409 | 1.0312 | 0.0298 | 0.9688 | -0.0564 | 0.0625 | 0.0217 |
| 1.0375 | 0.0572 | 0.9625 | -0.1305 | 0.0750 | 0.0488 | 1.0375 | 0.0337 | 0.9625 | -0.0730 | 0.0750 | 0.0319 |
| 1.0437 | 0.0642 | 0.9563 | -0.1674 | 0.0875 | 0.0601 | 1.0437 | 0.0376 | 0.9563 | -0.0895 | 0.0875 | 0.0342 |
| 1.0500 | 0.0721 | 0.9500 | -0.2024 | 0.1000 | 0.0681 | 1.0500 | 0.0415 | 0.9500 | -0.1051 | 0.1000 | 0.0422 |
| 1.0562 | 0.0791 | 0.9437 | -0.2453 | 0.1125 | 0.0817 | 1.0562 | 0.0454 | 0.9437 | -0.1363 | 0.1125 | 0.0468 |
| 1.0625 | 0.0869 | 0.9375 | -0.2959 | 0.1250 | 0.0964 | 1.0625 | 0.0486 | 0.9375 | -0.1596 | 0.1250 | 0.0558 |
| 1.0688 | 0.0940 | 0.9313 | -0.3543 | 0.1375 | 0.1133 | 1.0688 | 0.0533 | 0.9313 | -0.1946 | 0.1375 | 0.0627 |
| 1.0750 | 0.1050 | 0.9250 | -0.4127 | 0.1500 | 0.1347 | 1.0750 | 0.0580 | 0.9250 | -0.2297 | 0.1500 | 0.0751 |
| 1.0813 | 0.1151 | 0.9187 | -0.4827 | 0.1625 | 0.1596 | 1.0813 | 0.0634 | 0.9187 | -0.2764 | 0.1625 | 0.0853 |
| 1.0875 | 0.1292 | 0.9125 | -0.5723 | 0.1750 | 0.1878 | 1.0875 | 0.0697 | 0.9125 | -0.3270 | 0.1750 | 0.1011 |
| 1.0938 | 0.1418 | 0.9062 | -0.6657 | 0.1875 | 0.2227 | 1.0938 | 0.0775 | 0.9062 | -0.3854 | 0.1875 | 0.1192 |
| 1.1000 | 0.1582 | 0.9000 | -0.7591 | 0.2000 | 0.2611 | 1.1000 | 0.0862 | 0.9000 | -0.4555 | 0.2000 | 0.1429 |

## 3 Results

Table 3 and Figure 4 summarize and illustrate the discovered models for the human cortex for the tension, compression, and shear data from Table 1 using the isotropic, perfectly incompressible Constitutive Artificial Neural Network with two hidden layers from Figure 3. Table 3 summarizes the 24 weights *w*_1:2,1:12_ and the goodness of fit *R*^2^ for single-mode training with the three individual modes and for multi-mode training with all three modes combined. Figure 4 directly compares the data and model in terms of the nominal stress as a function of the stretch and shear strain, where the dots indicate the measured data and the color-coded regions highlight the individual contributions of the twelve network nodes to the discovered free energy function *ψ*. Warm red-type colors are associated with the first invariant, [*I*_1_ − 3] and cold blue-type colors are associated with the second invariant, [*I*_2_ − 3]. First, for single-mode training with the individual modes, the Constitutive Artificial Neural Network succeeds in *interpolating* or *fitting* the three individual sets of training data: The learned network parameters define stress functions that fit the individual tension, compression, and shear data excellently with 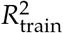 values of 0.99, 1.00, and 1.00. Second, for single-mode training, we observe that the network performs moderately at *extrapolating* or *predicting* data outside the training regime: The network parameters trained individually for each mode do not predict the other modes well, with 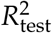 values ranging from 0.00 for the tension prediction with compression training to 0.93 for the shear prediction with tension training. Third, for multi-mode training with all data sets combined, we find that the goodness of fit 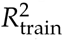 of the individual tests decreases to 0.36, 0.90, and 0.99, while the sum of the three *R*^2^ values, the collective fit of all three tests, increases. Fourth, for single-mode training, all twelve terms of the model are activated as indicated through the full color spectrum in the first three columns. At the same time, for for multi-mode training, the Constitutive Artificial Neural Network discovers a model with only four terms, while the weights of the other terms train to zero. Strikingly, these four terms are all functions of the second invariant, [*I*_2_ − 3], as indicated through the cold blue-type colors.

**Table 3:**
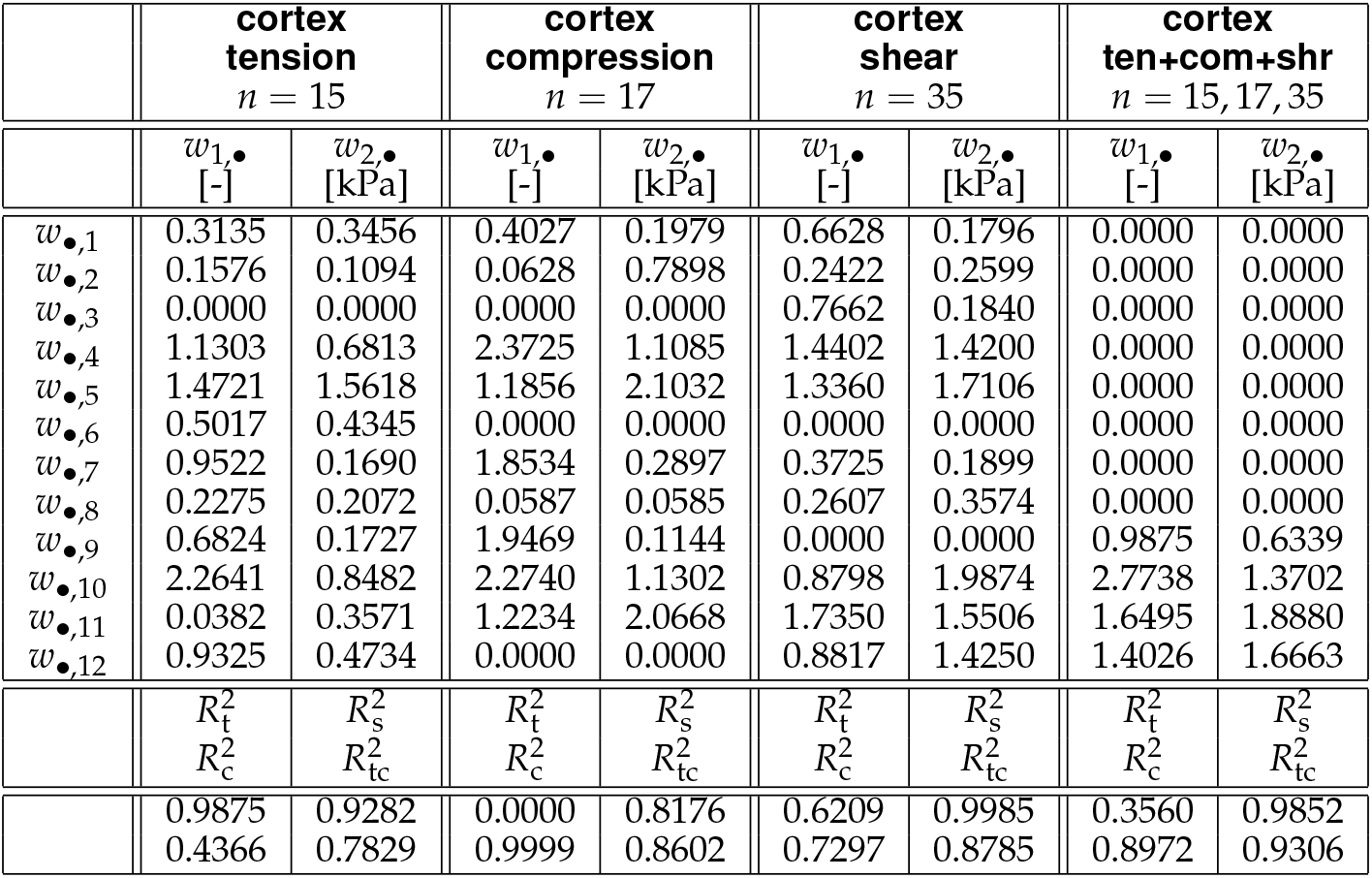
Gray matter model. Cortex parameters learned for tension, compression, and shear data from Table 1 using the isotropic, perfectly incompressible Constitutive Artificial Neural Network with two hidden layers, and twelve nodes from Figure 3. Summary of the 24 weights *w*_1:2,1:12_ and the goodness of fit *R*^2^ for training with the three individual tests and for all three tests combined.

| | cortex<br>tension<br>$n = 15$ | | cortex<br>compression<br>$n = 17$ | | cortex<br>shear<br>$n = 35$ | | cortex<br>ten+com+shr<br>$n = 15, 17, 35$ | |
| --- | --- | --- | --- | --- | --- | --- | --- | --- |
| | $w_{1,\bullet}$<br>[-] | $w_{2,\bullet}$<br>[kPa] | $w_{1,\bullet}$<br>[-] | $w_{2,\bullet}$<br>[kPa] | $w_{1,\bullet}$<br>[-] | $w_{2,\bullet}$<br>[kPa] | $w_{1,\bullet}$<br>[-] | $w_{2,\bullet}$<br>[kPa] |
| $w_{\bullet,1}$ | 0.3135 | 0.3456 | 0.4027 | 0.1979 | 0.6628 | 0.1796 | 0.0000 | 0.0000 |
| $w_{\bullet,2}$ | 0.1576 | 0.1094 | 0.0628 | 0.7898 | 0.2422 | 0.2599 | 0.0000 | 0.0000 |
| $w_{\bullet,3}$ | 0.0000 | 0.0000 | 0.0000 | 0.0000 | 0.7662 | 0.1840 | 0.0000 | 0.0000 |
| $w_{\bullet,4}$ | 1.1303 | 0.6813 | 2.3725 | 1.1085 | 1.4402 | 1.4200 | 0.0000 | 0.0000 |
| $w_{\bullet,5}$ | 1.4721 | 1.5618 | 1.1856 | 2.1032 | 1.3360 | 1.7106 | 0.0000 | 0.0000 |
| $w_{\bullet,6}$ | 0.5017 | 0.4345 | 0.0000 | 0.0000 | 0.0000 | 0.0000 | 0.0000 | 0.0000 |
| $w_{\bullet,7}$ | 0.9522 | 0.1690 | 1.8534 | 0.2897 | 0.3725 | 0.1899 | 0.0000 | 0.0000 |
| $w_{\bullet,8}$ | 0.2275 | 0.2072 | 0.0587 | 0.0585 | 0.2607 | 0.3574 | 0.0000 | 0.0000 |
| $w_{\bullet,9}$ | 0.6824 | 0.1727 | 1.9469 | 0.1144 | 0.0000 | 0.0000 | 0.9875 | 0.6339 |
| $w_{\bullet,10}$ | 2.2641 | 0.8482 | 2.2740 | 1.1302 | 0.8798 | 1.9874 | 2.7738 | 1.3702 |
| $w_{\bullet,11}$ | 0.0382 | 0.3571 | 1.2234 | 2.0668 | 1.7350 | 1.5506 | 1.6495 | 1.8880 |
| $w_{\bullet,12}$ | 0.9325 | 0.4734 | 0.0000 | 0.0000 | 0.8817 | 1.4250 | 1.4026 | 1.6663 |
| | $R_t^2$ | $R_s^2$ | $R_t^2$ | $R_s^2$ | $R_t^2$ | $R_s^2$ | $R_t^2$ | $R_s^2$ |
| | $R_c^2$ | $R_{tc}^2$ | $R_c^2$ | $R_{tc}^2$ | $R_c^2$ | $R_{tc}^2$ | $R_c^2$ | $R_{tc}^2$ |
|  | 0.9875 | 0.9282 | 0.0000 | 0.8176 | 0.6209 | 0.9985 | 0.3560 | 0.9852 |
|  | 0.4366 | 0.7829 | 0.9999 | 0.8602 | 0.7297 | 0.8785 | 0.8972 | 0.9306 |

**Figure 4:**
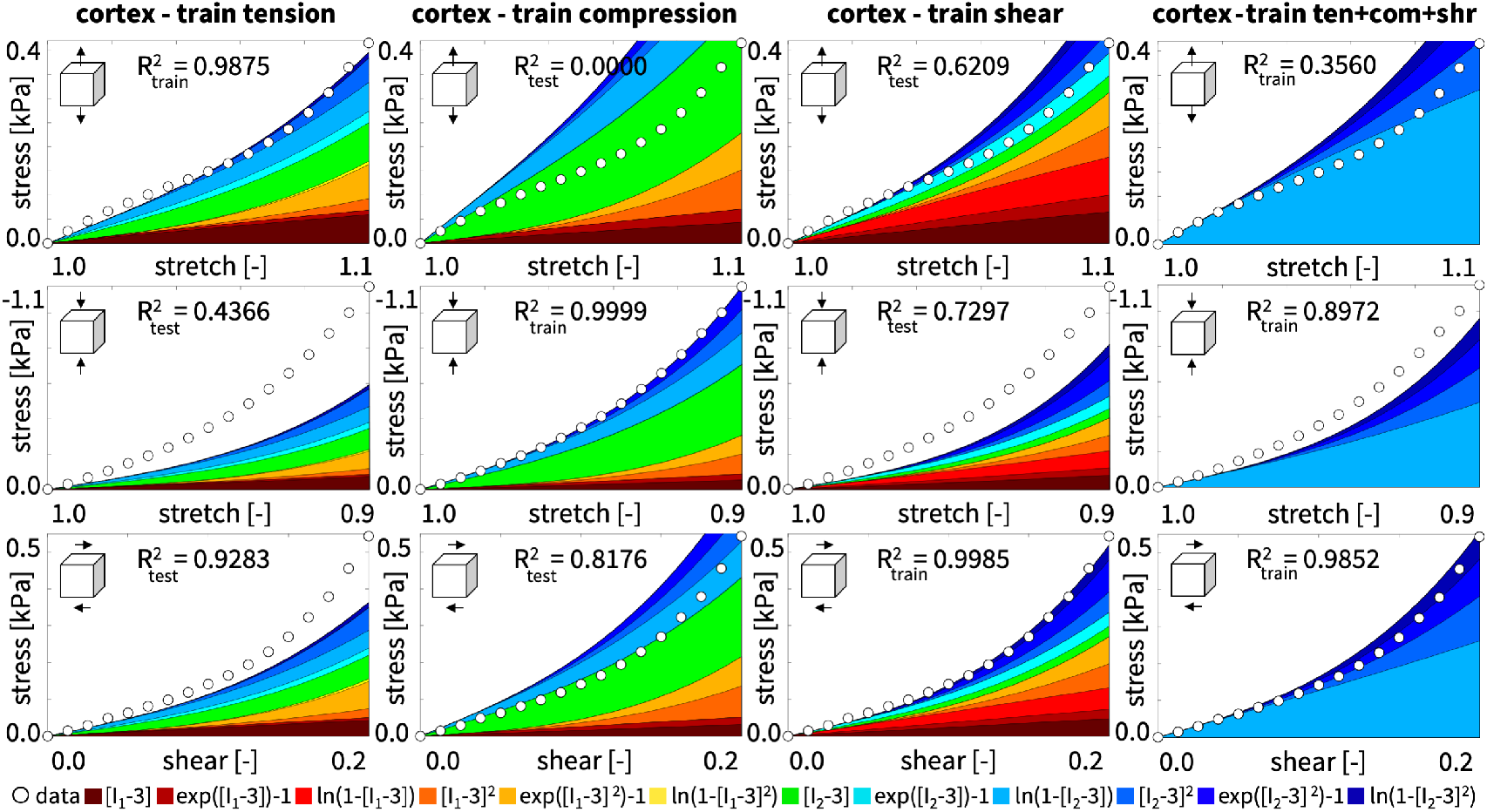
Gray matter data vs. model. Nominal stress as a function of stretch and shear strain for the isotropic, perfectly incompressible Constitutive Artificial Neural Network with two hidden layers, and twelve nodes from Figure 3. Dots illustrate the tension, compression, and shear data of the human cortex [11] from Table 1; color-coded areas highlight the twelve contributions to the discovered stress function according to Figure 3 from Table 3.

Table 4 and Figure 5 summarize and illustrate the discovered models for the human corona radiata for the tension, compression, and shear data from Table 2. The white matter results from the corona radiata confirm the trends of the gray matter results for the cortex in Table 3 and Figure 4. First, for single-mode training, our neural network succeeds in *interpolating* or *fitting* the individual training data: The learned network parameters define stress functions that fit the individual tension, compression, and shear data excellently with 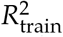 values of 0.99, 1.00, and 1.00. Second, the network performs moderately at *extrapolating* or *predicting* data outside the training regime: The network parameters trained for each individual mode fail to predict the other modes equally well, with 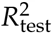 values ranging from 0.0 for the tension predictions with both compression and shear training to 0.83 for the shear prediction with tension training. Third, we find that, for all tests combined, the goodness of fit 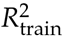 of the tensile test remains 0.00 and decreases to 0.79 and 0.93 for compression and shear, but the collective fit increases. Fourth, similar to the gray matter results in Figure 4, the white matter model trained with the individual tests in Figure 5 activates seven, eight, and eleven terms as indicated through the broad color spectrum in the first three columns. Interestingly, for all three tests combined, the Constitutive Artificial Neural Network discovers a model with only four terms, which are again all functions of the second invariant, [*I*_2_ − 3], as indicated through the cold blue-type colors.

**Table 4:** White matter model. Corona radiata parameters learned for tension, compression, and shear data from Table 2 using the isotropic, perfectly incompressible Constitutive Artificial Neural Network with two hidden layers, and twelve nodes from Figure 3. Summary of the 24 weights *w*_1:2,1:12_ and the goodness of fit *R*^2^ for training with the three individual tests and with all three tests combined.

| | corona radiata<br>tension<br>$n = 18$ | | corona radiata<br>compression<br>$n = 18$ | | corona radiata<br>shear<br>$n = 33$ | | corona radiata<br>ten+com+shr<br>$n = 18, 18, 33$ | |
| --- | --- | --- | --- | --- | --- | --- | --- | --- |
| | $w_{1,\bullet}$<br>[-] | $w_{2,\bullet}$<br>[kPa] | $w_{1,\bullet}$<br>[-] | $w_{2,\bullet}$<br>[kPa] | $w_{1,\bullet}$<br>[-] | $w_{2,\bullet}$<br>[kPa] | $w_{1,\bullet}$<br>[-] | $w_{2,\bullet}$<br>[kPa] |
| $w_{\bullet,1}$ | 0.0000 | 0.0000 | 1.7357 | 0.2807 | 0.3643 | 0.2492 | 0.0000 | 0.0000 |
| $w_{\bullet,2}$ | 0.0000 | 0.0000 | 0.0000 | 0.0000 | 0.1032 | 0.2404 | 0.0000 | 0.0000 |
| $w_{\bullet,3}$ | 0.0000 | 0.0000 | 0.0000 | 0.0000 | 0.0369 | 0.3070 | 0.0000 | 0.0000 |
| $w_{\bullet,4}$ | 0.8932 | 0.1474 | 1.5473 | 1.0767 | 1.3942 | 0.6520 | 0.0000 | 0.0000 |
| $w_{\bullet,5}$ | 0.3760 | 0.2325 | 1.1415 | 1.2150 | 1.3600 | 1.1027 | 0.0000 | 0.0000 |
| $w_{\bullet,6}$ | 1.3081 | 0.4295 | 1.2115 | 1.1480 | 0.4401 | 0.8310 | 0.0000 | 0.0000 |
| $w_{\bullet,7}$ | 1.0042 | 0.0717 | 0.0000 | 0.0000 | 0.0349 | 0.2945 | 1.3862 | 0.1598 |
| $w_{\bullet,8}$ | 0.0867 | 0.0717 | 0.0029 | 0.0295 | 0.0550 | 0.3905 | 0.2398 | 0.4900 |
| $w_{\bullet,9}$ | 0.8403 | 0.2065 | 0.0000 | 0.0000 | 0.7680 | 0.1179 | 0.0000 | 0.0000 |
| $w_{\bullet,10}$ | 0.0000 | 0.0000 | 1.0083 | 1.4130 | 1.0552 | 0.8552 | 0.0000 | 0.0000 |
| $w_{\bullet,11}$ | 0.0000 | 0.0000 | 1.2191 | 1.1331 | 0.9990 | 1.0740 | 1.8893 | 1.6859 |
| $w_{\bullet,12}$ | 1.1048 | 0.0030 | 2.6478 | 0.8233 | 0.0000 | 0.0000 | 1.1789 | 1.9113 |
| | $R_t^2$ | $R_s^2$ | $R_t^2$ | $R_s^2$ | $R_t^2$ | $R_s^2$ | $R_t^2$ | $R_s^2$ |
| | $R_c^2$ | $R_{tc}^2$ | $R_c^2$ | $R_{tc}^2$ | $R_c^2$ | $R_{tc}^2$ | $R_c^2$ | $R_{tc}^2$ |
|  | 0.9875 | 0.8250 | 0.0000 | 0.0620 | 0.0000 | 0.9989 | 0.0000 | 0.9349 |
|  | 0.0460 | 0.5471 | 0.9998 | 0.6251 | 0.4837 | 0.7271 | 0.7898 | 0.8361 |

**Figure 5:**
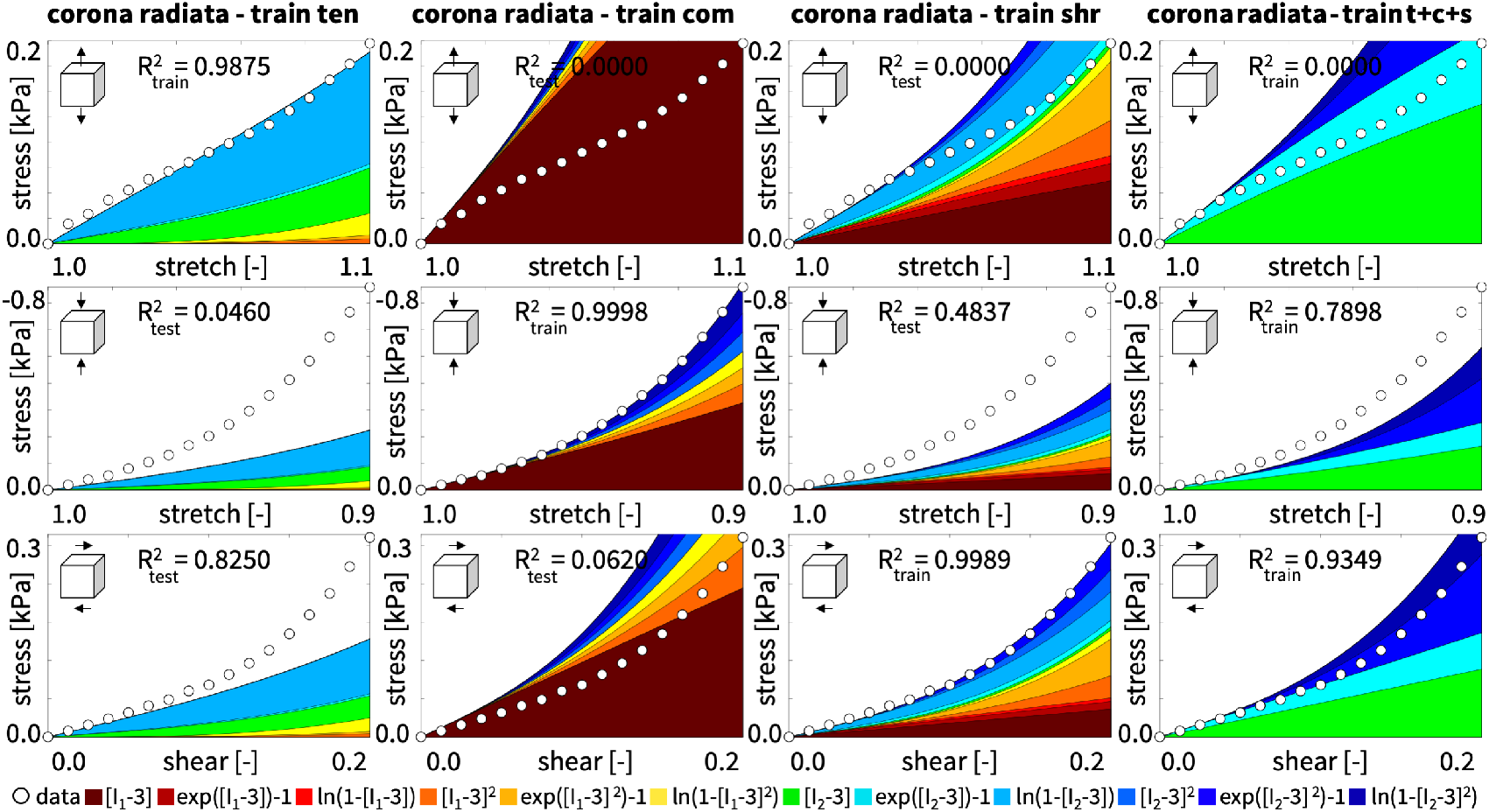
White matter data vs. model. Nominal stress as a function of stretch and shear strain for the isotropic, perfectly incompressible Constitutive Artificial Neural Network with two hidden layers, and twelve nodes from Figure 3. Dots illustrate the tension, compression, and shear data of the human corona radiata [11] from Table 1; color-coded areas highlight the twelve contributions to the discovered stress function according to Figure 3 from Table 4.

Table 5 and Figure 6 summarize and illustrate the discovered models for the human cortex, basal ganglia, corona radiata, and corpus callosum for multi-mode training with the tension, compression, and shear data from Tables 1 and 2. First and foremost, for multi-mode training, the fit of the shear data with 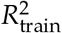 values of 0.99, 0.97, 0.93, and 0.92 is uniformly the best across all four brain regions. Second, the model universally *underestimates* the compressive stresses with *R*^*2*^ values ranging from 0.78 to 0.90, and *overestimates* the tensile stresses with *R*^2^ values from 0.00 to 0.36, indicating a poor fit of the tensile data. Third, and most importantly, the side-by-side comparison of all four brain regions confirms the trends of the cortex and the corona radiata: Our Constitutive Artificial Neural Network uniquely discovers a family of models that is parameterized in terms of the *second invariant only*, while the weights of the first invariant terms consistently train to zero. The blue color spectrum in Figure 6 underscores this observation.

**Table 5:** Gray and white matter models. Cortex, basal ganglia, corona radiata, and corpus callosum parameters learned for combined tension, compression, and shear data from Tables 1 and 2 using the isotropic, perfectly incompressible Constitutive Artificial Neural Network with two hidden layers, and twelve nodes from Figure 3. Summary of the 12 non-zero weights *w*_1:2,7:12_ and the goodness of fit *R*^2^ for training with all three tests combined.

| | **cortex**<br>ten+com+shr<br>$n = 15, 17, 35$ | | **basal ganglia**<br>ten+com+shr<br>$n = 15, 15, 29$ | | **corona radiata**<br>ten+com+shr<br>$n = 18, 18, 36$ | | **corpus callosum**<br>ten+com+shr<br>$n = 19, 20, 39$ | |
| --- | --- | --- | --- | --- | --- | --- | --- | --- |
| | $w_{1,\bullet}$<br>[-] | $w_{2,\bullet}$<br>[kPa] | $w_{1,\bullet}$<br>[-] | $w_{2,\bullet}$<br>[kPa] | $w_{1,\bullet}$<br>[-] | $w_{2,\bullet}$<br>[kPa] | $w_{1,\bullet}$<br>[-] | $w_{2,\bullet}$<br>[kPa] |
| $w_{\bullet,7}$ | 0.0000 | 0.0000 | 1.7880 | 0.1927 | 1.3862 | 0.1598 | 0.5635 | 0.1067 |
| $w_{\bullet,8}$ | 0.0000 | 0.0000 | 0.0000 | 0.0000 | 0.2398 | 0.4900 | 0.2363 | 0.1383 |
| $w_{\bullet,9}$ | 0.9875 | 0.6339 | 0.0000 | 0.0000 | 0.0000 | 0.0000 | 0.8398 | 0.1135 |
| $w_{\bullet,10}$ | 2.7738 | 1.3702 | 0.9396 | 0.8143 | 0.0000 | 0.0000 | 0.9210 | 1.1218 |
| $w_{\bullet,11}$ | 1.6495 | 1.8880 | 1.6193 | 1.1867 | 1.8893 | 1.6859 | 1.0628 | 1.0185 |
| $w_{\bullet,12}$ | 1.4026 | 1.6663 | 0.9666 | 0.7932 | 1.1789 | 1.9113 | 0.7282 | 1.2621 |
| | $R^2_t$<br>$R^2_c$ | $R^2_s$<br>$R^2_{tc}$ | $R^2_t$<br>$R^2_c$ | $R^2_s$<br>$R^2_{tc}$ | $R^2_t$<br>$R^2_c$ | $R^2_s$<br>$R^2_{tc}$ | $R^2_t$<br>$R^2_c$ | $R^2_s$<br>$R^2_{tc}$ |
|  | 0.3560<br>0.8972 | 0.9852<br>0.9306 | 0.0000<br>0.8646 | 0.9739<br>0.9135 | 0.0000<br>0.7898 | 0.9349<br>0.8361 | 0.0000<br>0.7847 | 0.9209<br>0.8303 |

**Figure 6:**
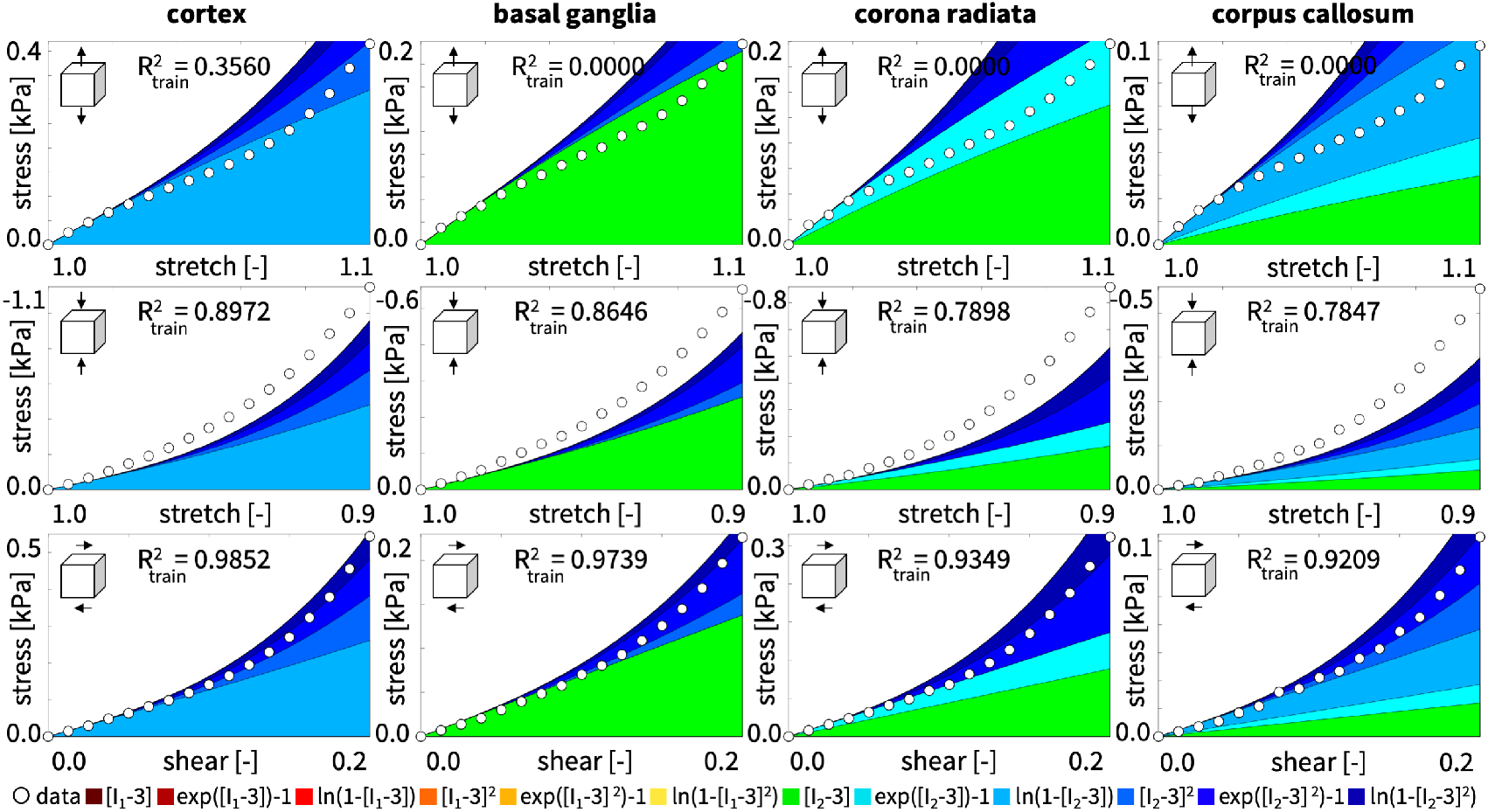
Gray and white matter data vs. model. Nominal stress as a function of stretch and shear strain for the isotropic, perfectly incompressible Constitutive Artificial Neural Network with two hidden layers, and twelve nodes from Figure 3. Dots illustrate the tension, compression, and shear data of all four brain regions [11] from Table 1; color-coded areas highlight the six contributions to the discovered stress function according to Figure 3 from Table 5.

Table 6 and Figure 7 highlight the effects of the L2 regularization according to equation (23). As expected, the regularization reduces the number of non-zero terms, in our case from six in Table 5 and Figure 6, to two for the cortex, the corona radiata, and the corpus callosum, and only one for the basal ganglia. The associated non-zero weights, *w*_1,8_, *w*_2,8_, *w*_1,9_, *w*_2,9_, activate the linear exponential term, exp([I_2_ − 3]) − 1, and the linear logarithmic term, ln(1 − [I_2_ − 3]), which are highlighted in turquoise and light blue in Figure 7. The general trends are the same for the discovered six-term model without regularization and two-term model with L2 regularization: Both models depend on the second invariant only and their fits are best for the shear data with *R*^2^ values well above 0.90 and worst for the tension data with *R*^2^ values ranging from 0.00 to 0.48.

**Table 6:** Gray and white matter models. Cortex, basal ganglia, corona radiata, and corpus callosum parameters learned for combined tension, compression, and shear data from Tables 1 and 2 using the isotropic, perfectly incompressible Constitutive Artificial Neural Network with two hidden layers, and twelve nodes from Figure 3 with additional L2 regularization for the weights. Summary of the four non-zero weights *w*_1:2,8:9_ and the goodness of fit *R*^2^ for training with all three tests combined.

| | cortex<br>ten+com+shr<br>$n = 15, 17, 35$ | | basal ganglia<br>ten+com+shr<br>$n = 15, 15, 29$ | | corona radiata<br>ten+com+shr<br>$n = 18, 18, 36$ | | corpus callosum<br>ten+com+shr<br>$n = 19, 20, 39$ | |
| --- | --- | --- | --- | --- | --- | --- | --- | --- |
| | $w_{1,\bullet}$<br>[-] | $w_{2,\bullet}$<br>[kPa] | $w_{1,\bullet}$<br>[-] | $w_{2,\bullet}$<br>[kPa] | $w_{1,\bullet}$<br>[-] | $w_{2,\bullet}$<br>[kPa] | $w_{1,\bullet}$<br>[-] | $w_{2,\bullet}$<br>[kPa] |
| $w_{\bullet,8}$ | 0.4957 | 0.4442 | 0.0000 | 0.0000 | 0.4560 | 0.4351 | 0.2409 | 0.2367 |
| $w_{\bullet,9}$ | 0.9840 | 0.7064 | 0.6802 | 0.6250 | 0.5614 | 0.4987 | 0.4739 | 0.4551 |
| | $R_t^2$ | $R_s^2$ | $R_t^2$ | $R_s^2$ | $R_t^2$ | $R_s^2$ | $R_t^2$ | $R_s^2$ |
| | $R_c^2$ | $R_{tc}^2$ | $R_c^2$ | $R_{tc}^2$ | $R_c^2$ | $R_{tc}^2$ | $R_c^2$ | $R_{tc}^2$ |
|  | 0.4590 | 0.9477 | 0.4778 | 0.9699 | 0.0000 | 0.9551 | 0.0000 | 0.9620 |
|  | 0.7425 | 0.8788 | 0.6696 | 0.8588 | 0.5143 | 0.7418 | 0.5109 | 0.7276 |

**Figure 7:**
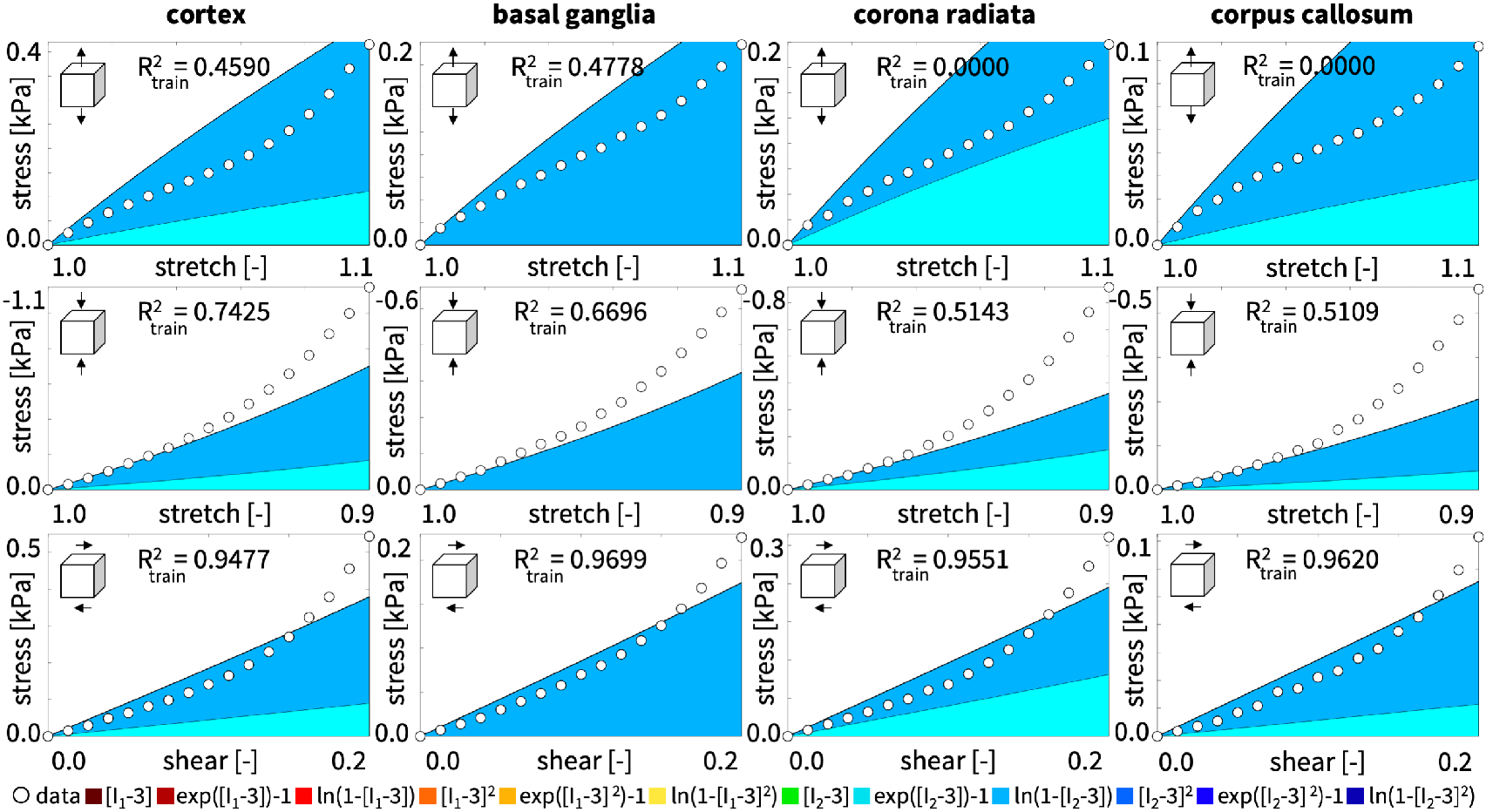
Gray and white matter data vs. model. Nominal stress as a function of stretch and shear strain for the isotropic, perfectly incompressible Constitutive Artificial Neural Network with two hidden layers, and twelve nodes from Figure 3 with additional L2 regularization for the weights. Dots illustrate the tension, compression, and shear data of all four brain regions [11] from Table 1; color-coded areas highlight the two contributions to the discovered stress function according to Figure 3 from Table 6.

Table 7 and Figure 8 demonstrate an application of our Constitutive Artificial Neural Network beyond model discovery, the parameter identification and comparison of special cases of the generalized network according to equations (16) to (21). The first set of models, the neo Hooke, Blatz Ko, and Mooney Rivlin models, are all linear in terms of the first invariant, second invariant, or both; the second set, the Demiray, Gent, and Holzapfel models, contain linear exponential, linear logarithmic, or quadratic exponential terms. Table 7 shows that each model, except for the Mooney Rivlin model, activates only one term of our network, either the first, second, third, fifth, or seventh. For all six models, we can convert the weights into a stiffness-like parameter with units [kPa]; the linear Mooney Rivlin model has an additional stiffness-like parameter, and the three nonlinear models have an additional coefficient of nonlinearity. Figure 8 shows the behavior of the neo Hooke, Blatz Ko, Demiray, and Holzapfel models when simultaneously trained for the tension, compression, and shear data of the human cortex. Notably, for the small stretch and shear strain ranges of 0.9 ≤ *λ* ≤ 1.1 and 0.0 ≤ *γ* ≤ 0.2, only the Holzapfel model displays a marked strain stiffening, while the neo Hooke, Blatz Ko, and Demiray models remain is their predominantly linear regimes. This allows the Holzapfel model to perform best not only in shear, with an *R*^2^ values of 0.96, but also in tension with a value of 0.48, where most other models fail.

**Table 7:**
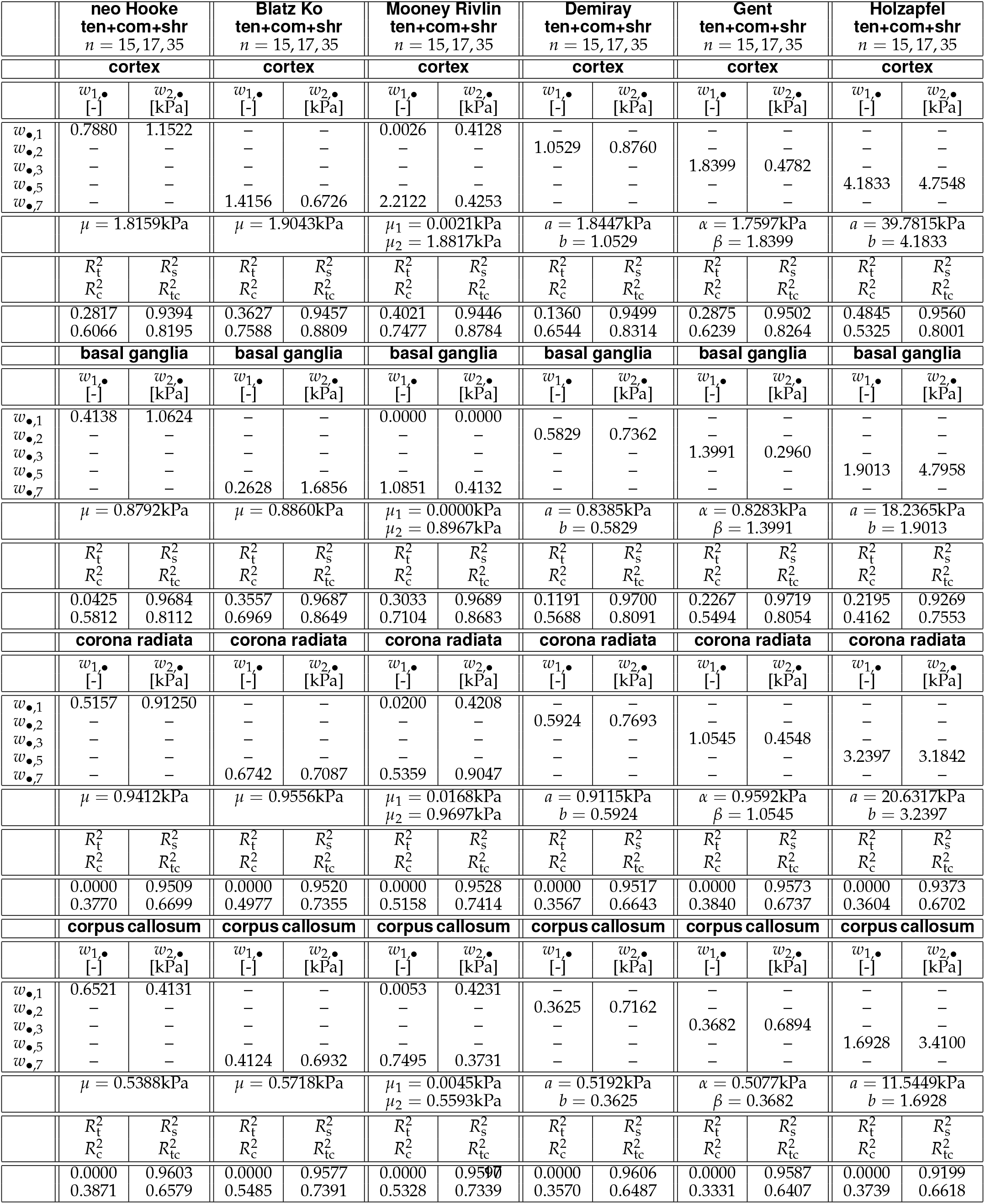
Special cases of neo Hooke, Blatz Ko, Mooney Rivlin, Demiray, Gent, and Holzapfel models. Cortex, basal ganglia, corona radiata, and corpus callosum parameters learned for combined tension, compression, and shear data from Tables 1 and 2. Summary of the non-zero weights, the physics parameters *μ, μ*_1_, *μ*_2_, *a, b, α, β*, and the goodness of fit *R*^2^ for training with all three tests combined.

| | **neo Hooke**<br>**ten+com+shr**<br>$n = 15, 17, 35$ | | **Blatz Ko**<br>**ten+com+shr**<br>$n = 15, 17, 35$ | | **Mooney Rivlin**<br>**ten+com+shr**<br>$n = 15, 17, 35$ | | **Demiray**<br>**ten+com+shr**<br>$n = 15, 17, 35$ | | **Gent**<br>**ten+com+shr**<br>$n = 15, 17, 35$ | | **Holzapfel**<br>**ten+com+shr**<br>$n = 15, 17, 35$ | |
| --- | --- | --- | --- | --- | --- | --- | --- | --- | --- | --- | --- | --- |
|  | **cortex** |  | **cortex** |  | **cortex** |  | **cortex** |  | **cortex** |  | **cortex** |  |
| | $w_{1,\bullet}$<br>[-] | $w_{2,\bullet}$<br>[kPa] | $w_{1,\bullet}$<br>[-] | $w_{2,\bullet}$<br>[kPa] | $w_{1,\bullet}$<br>[-] | $w_{2,\bullet}$<br>[kPa] | $w_{1,\bullet}$<br>[-] | $w_{2,\bullet}$<br>[kPa] | $w_{1,\bullet}$<br>[-] | $w_{2,\bullet}$<br>[kPa] | $w_{1,\bullet}$<br>[-] | $w_{2,\bullet}$<br>[kPa] |
| $w_{\bullet,1}$ | 0.7880 | 1.1522 | — | — | 0.0026 | 0.4128 | — | — | — | — | — | — |
| $w_{\bullet,2}$ | — | — | — | — | — | — | 1.0529 | 0.8760 | — | — | — | — |
| $w_{\bullet,3}$ | — | — | — | — | — | — | — | — | 1.8399 | 0.4782 | — | — |
| $w_{\bullet,5}$ | — | — | — | — | — | — | — | — | — | — | 4.1833 | 4.7548 |
| $w_{\bullet,7}$ | — | — | 1.4156 | 0.6726 | 2.2122 | 0.4253 | — | — | — | — | — | — |
| | $\mu = 1.8159\text{kPa}$ | | $\mu = 1.9043\text{kPa}$ | | $\mu_1 = 0.0021\text{kPa}$<br>$\mu_2 = 1.8817\text{kPa}$ | | $a = 1.8447\text{kPa}$<br>$b = 1.0529$ | | $\alpha = 1.7597\text{kPa}$<br>$\beta = 1.8399$ | | $a = 39.7815\text{kPa}$<br>$b = 4.1833$ | |
| | $R_t^2$<br>$R_c^2$ | $R_s^2$<br>$R_{tc}^2$ | $R_t^2$<br>$R_c^2$ | $R_s^2$<br>$R_{tc}^2$ | $R_t^2$<br>$R_c^2$ | $R_s^2$<br>$R_{tc}^2$ | $R_t^2$<br>$R_c^2$ | $R_s^2$<br>$R_{tc}^2$ | $R_t^2$<br>$R_c^2$ | $R_s^2$<br>$R_{tc}^2$ | $R_t^2$<br>$R_c^2$ | $R_s^2$<br>$R_{tc}^2$ |
|  | 0.2817<br>0.6066 | 0.9394<br>0.8195 | 0.3627<br>0.7588 | 0.9457<br>0.8809 | 0.4021<br>0.7477 | 0.9446<br>0.8784 | 0.1360<br>0.6544 | 0.9499<br>0.8314 | 0.2875<br>0.6239 | 0.9502<br>0.8264 | 0.4845<br>0.5325 | 0.9560<br>0.8001 |
|  | **basal ganglia** |  | **basal ganglia** |  | **basal ganglia** |  | **basal ganglia** |  | **basal ganglia** |  | **basal ganglia** |  |
| | $w_{1,\bullet}$<br>[-] | $w_{2,\bullet}$<br>[kPa] | $w_{1,\bullet}$<br>[-] | $w_{2,\bullet}$<br>[kPa] | $w_{1,\bullet}$<br>[-] | $w_{2,\bullet}$<br>[kPa] | $w_{1,\bullet}$<br>[-] | $w_{2,\bullet}$<br>[kPa] | $w_{1,\bullet}$<br>[-] | $w_{2,\bullet}$<br>[kPa] | $w_{1,\bullet}$<br>[-] | $w_{2,\bullet}$<br>[kPa] |
| $w_{\bullet,1}$ | 0.4138 | 1.0624 | — | — | 0.0000 | 0.0000 | — | — | — | — | — | — |
| $w_{\bullet,2}$ | — | — | — | — | — | — | 0.5829 | 0.7362 | — | — | — | — |
| $w_{\bullet,3}$ | — | — | — | — | — | — | — | — | 1.3991 | 0.2960 | — | — |
| $w_{\bullet,5}$ | — | — | — | — | — | — | — | — | — | — | 1.9013 | 4.7958 |
| $w_{\bullet,7}$ | — | — | 0.2628 | 1.6856 | 1.0851 | 0.4132 | — | — | — | — | — | — |
| | $\mu = 0.8792\text{kPa}$ | | $\mu = 0.8860\text{kPa}$ | | $\mu_1 = 0.0000\text{kPa}$<br>$\mu_2 = 0.8967\text{kPa}$ | | $a = 0.8385\text{kPa}$<br>$b = 0.5829$ | | $\alpha = 0.8283\text{kPa}$<br>$\beta = 1.3991$ | | $a = 18.2365\text{kPa}$<br>$b = 1.9013$ | |
| | $R_t^2$<br>$R_c^2$ | $R_s^2$<br>$R_{tc}^2$ | $R_t^2$<br>$R_c^2$ | $R_s^2$<br>$R_{tc}^2$ | $R_t^2$<br>$R_c^2$ | $R_s^2$<br>$R_{tc}^2$ | $R_t^2$<br>$R_c^2$ | $R_s^2$<br>$R_{tc}^2$ | $R_t^2$<br>$R_c^2$ | $R_s^2$<br>$R_{tc}^2$ | $R_t^2$<br>$R_c^2$ | $R_s^2$<br>$R_{tc}^2$ |
|  | 0.0425<br>0.5812 | 0.9684<br>0.8112 | 0.3557<br>0.6969 | 0.9687<br>0.8649 | 0.3033<br>0.7104 | 0.9689<br>0.8683 | 0.1191<br>0.5688 | 0.9700<br>0.8091 | 0.2267<br>0.5494 | 0.9719<br>0.8054 | 0.2195<br>0.4162 | 0.9269<br>0.7553 |
|  | **corona radiata** |  | **corona radiata** |  | **corona radiata** |  | **corona radiata** |  | **corona radiata** |  | **corona radiata** |  |
| | $w_{1,\bullet}$<br>[-] | $w_{2,\bullet}$<br>[kPa] | $w_{1,\bullet}$<br>[-] | $w_{2,\bullet}$<br>[kPa] | $w_{1,\bullet}$<br>[-] | $w_{2,\bullet}$<br>[kPa] | $w_{1,\bullet}$<br>[-] | $w_{2,\bullet}$<br>[kPa] | $w_{1,\bullet}$<br>[-] | $w_{2,\bullet}$<br>[kPa] | $w_{1,\bullet}$<br>[-] | $w_{2,\bullet}$<br>[kPa] |
| $w_{\bullet,1}$ | 0.5157 | 0.91250 | — | — | 0.0200 | 0.4208 | — | — | — | — | — | — |
| $w_{\bullet,2}$ | — | — | — | — | — | — | 0.5924 | 0.7693 | — | — | — | — |
| $w_{\bullet,3}$ | — | — | — | — | — | — | — | — | 1.0545 | 0.4548 | — | — |
| $w_{\bullet,5}$ | — | — | — | — | — | — | — | — | — | — | 3.2397 | 3.1842 |
| $w_{\bullet,7}$ | — | — | 0.6742 | 0.7087 | 0.5359 | 0.9047 | — | — | — | — | — | — |
| | $\mu = 0.9412\text{kPa}$ | | $\mu = 0.9556\text{kPa}$ | | $\mu_1 = 0.0168\text{kPa}$<br>$\mu_2 = 0.9697\text{kPa}$ | | $a = 0.9115\text{kPa}$<br>$b = 0.5924$ | | $\alpha = 0.9592\text{kPa}$<br>$\beta = 1.0545$ | | $a = 20.6317\text{kPa}$<br>$b = 3.2397$ | |
| | $R_t^2$<br>$R_c^2$ | $R_s^2$<br>$R_{tc}^2$ | $R_t^2$<br>$R_c^2$ | $R_s^2$<br>$R_{tc}^2$ | $R_t^2$<br>$R_c^2$ | $R_s^2$<br>$R_{tc}^2$ | $R_t^2$<br>$R_c^2$ | $R_s^2$<br>$R_{tc}^2$ | $R_t^2$<br>$R_c^2$ | $R_s^2$<br>$R_{tc}^2$ | $R_t^2$<br>$R_c^2$ | $R_s^2$<br>$R_{tc}^2$ |
|  | 0.0000<br>0.3770 | 0.9509<br>0.6699 | 0.0000<br>0.4977 | 0.9520<br>0.7355 | 0.0000<br>0.5158 | 0.9528<br>0.7414 | 0.0000<br>0.3567 | 0.9517<br>0.6643 | 0.0000<br>0.3840 | 0.9573<br>0.6737 | 0.0000<br>0.3604 | 0.9373<br>0.6702 |
|  | **corpus callosum** |  | **corpus callosum** |  | **corpus callosum** |  | **corpus callosum** |  | **corpus callosum** |  | **corpus callosum** |  |
| | $w_{1,\bullet}$<br>[-] | $w_{2,\bullet}$<br>[kPa] | $w_{1,\bullet}$<br>[-] | $w_{2,\bullet}$<br>[kPa] | $w_{1,\bullet}$<br>[-] | $w_{2,\bullet}$<br>[kPa] | $w_{1,\bullet}$<br>[-] | $w_{2,\bullet}$<br>[kPa] | $w_{1,\bullet}$<br>[-] | $w_{2,\bullet}$<br>[kPa] | $w_{1,\bullet}$<br>[-] | $w_{2,\bullet}$<br>[kPa] |
| $w_{\bullet,1}$ | 0.6521 | 0.4131 | — | — | 0.0053 | 0.4231 | — | — | — | — | — | — |
| $w_{\bullet,2}$ | — | — | — | — | — | — | 0.3625 | 0.7162 | — | — | — | — |
| $w_{\bullet,3}$ | — | — | — | — | — | — | — | — | 0.3682 | 0.6894 | — | — |
| $w_{\bullet,5}$ | — | — | — | — | — | — | — | — | — | — | 1.6928 | 3.4100 |
| $w_{\bullet,7}$ | — | — | 0.4124 | 0.6932 | 0.7495 | 0.3731 | — | — | — | — | — | — |
| | $\mu = 0.5388\text{kPa}$ | | $\mu = 0.5718\text{kPa}$ | | $\mu_1 = 0.0045\text{kPa}$<br>$\mu_2 = 0.5593\text{kPa}$ | | $a = 0.5192\text{kPa}$<br>$b = 0.3625$ | | $\alpha = 0.5077\text{kPa}$<br>$\beta = 0.3682$ | | $a = 11.5449\text{kPa}$<br>$b = 1.6928$ | |
| | $R_t^2$<br>$R_c^2$ | $R_s^2$<br>$R_{tc}^2$ | $R_t^2$<br>$R_c^2$ | $R_s^2$<br>$R_{tc}^2$ | $R_t^2$<br>$R_c^2$ | $R_s^2$<br>$R_{tc}^2$ | $R_t^2$<br>$R_c^2$ | $R_s^2$<br>$R_{tc}^2$ | $R_t^2$<br>$R_c^2$ | $R_s^2$<br>$R_{tc}^2$ | $R_t^2$<br>$R_c^2$ | $R_s^2$<br>$R_{tc}^2$ |
|  | 0.0000<br>0.3871 | 0.9603<br>0.6579 | 0.0000<br>0.5485 | 0.9577<br>0.7391 | 0.0000<br>0.5328 | 0.9590<br>0.7339 | 0.0000<br>0.3570 | 0.9606<br>0.6487 | 0.0000<br>0.3331 | 0.9587<br>0.6407 | 0.0000<br>0.3739 | 0.9199<br>0.6618 |

**Figure 8:**
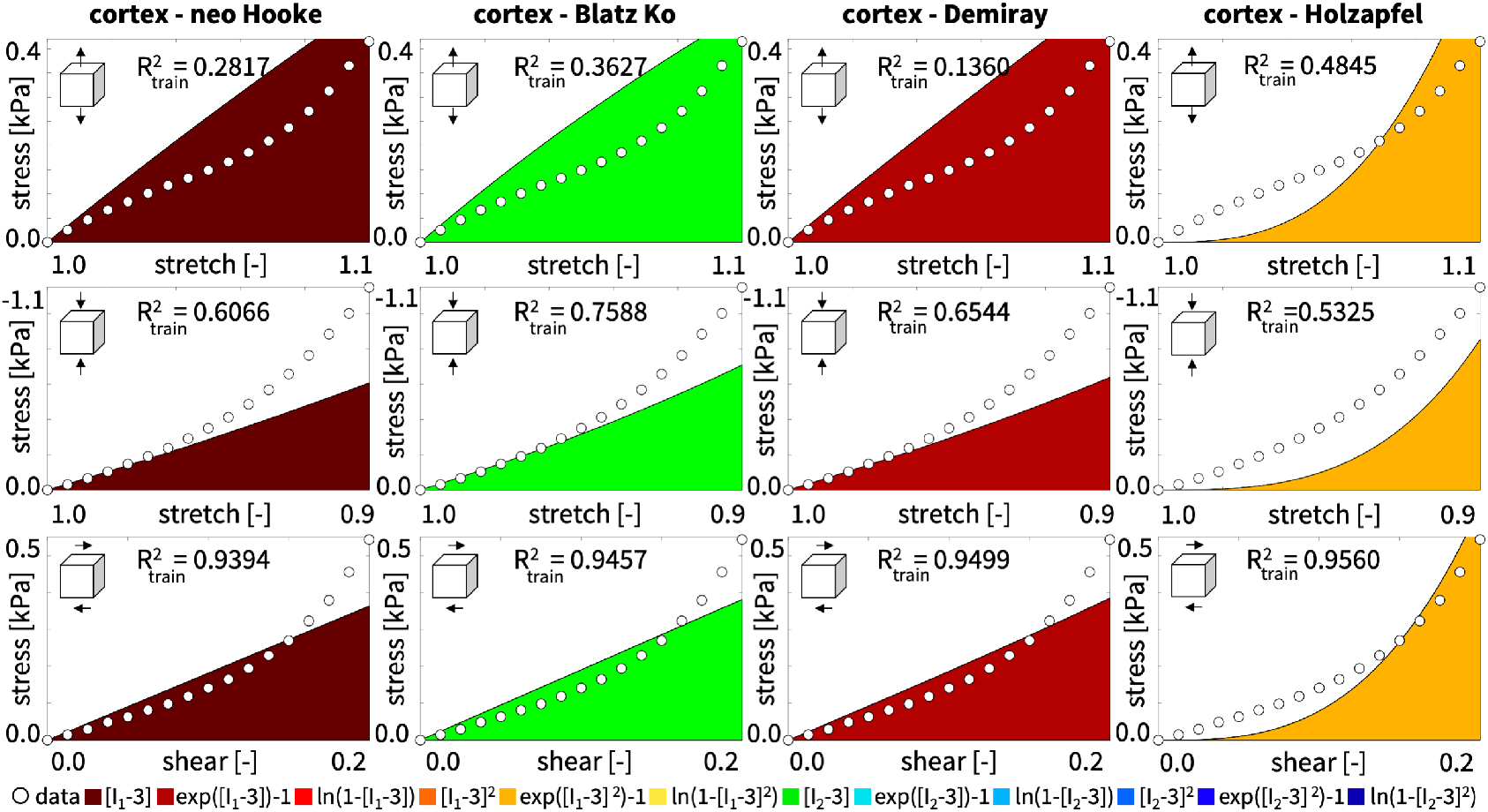
Special cases of neo Hooke, Blatz Ko, Mooney Rivlin, Demiray, Gent, and Holzapfel models. Nominal stress as a function of stretch and shear for special cases of the isotropic, perfectly incompressible Constitutive Artificial Neural Network from Figure 3. Dots illustrate the tension, compression, and shear data of the human cortex [11] from Table 1; color-coded areas highlight the terms of the stress function according to Figure 3 from Table 7.

Figure 9 summarizes and compares the performance of all models, the Constitutive Artificial Neural Network without and with L2 regularization and its special cases, the neo Hooke, Blatz Ko, Demiray, Gent, and Holzapfel models. The graphs in the first three columns result from single-mode training with the individual tension, compression, and shear data [11] from Tables 1 and 2; the last column results from multi-mode training with all three data sets combined. The three rows highlight the coefficients of determination for tension 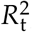, compression 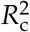, and shear 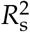. The color-coded blocks and error bars represent the means and standard deviations of the *R*^*2*^ value from 50 training runs with varying with varying random weight initializations. First, in the three graphs on the diagonal that reflect the *training* of the models, the 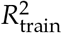 values of all seven models are close to one, with only three models training poorly, the neo Hooke and Holzapfel models in tension and the Blatz Ko model in compression. Notably, our non-regularized Constitutive Artificial Neural Network outperforms all other models and has the largest 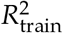 values when trained individually for tension, compression, and shear. Second, from the six off-diagonal graphs that reflect the *testing* of the models, we conclude that the model and parameters trained for tension are generally incapable of predicting the compression behavior and vice versa. However, the tension parameters are reasonably well suited to characterize the shear behavior, with our Constitutive Artificial Neural Network and the Holzapfel model performing best; vice versa, the shear parameters are moderately suited to characterize the tensile behavior, with our Constitutive Artificial Neural Network and the Blatz Ko model performing best. Finally, from the right column that reflects *training* with all three data sets combined, we conclude that our Constitutive Artificial Neural Network performs best for all three modes, followed by its L2 regularized counterpart, and the Blatz Ko model. Interestingly, we observe large *R*^2^ values across the entire bottom row, indicating that, of all three tests, the shear tests are generally the easiest to fit and predict for all seven models. Taken together, our non-regularized Constitutive Artificial Neural Network performs best in eight of all twelve cases, second best in two, and fifths in one suggesting that our proposed neural network *successfully discovers both model and parameters* that best describe the data.

**Figure 9:**
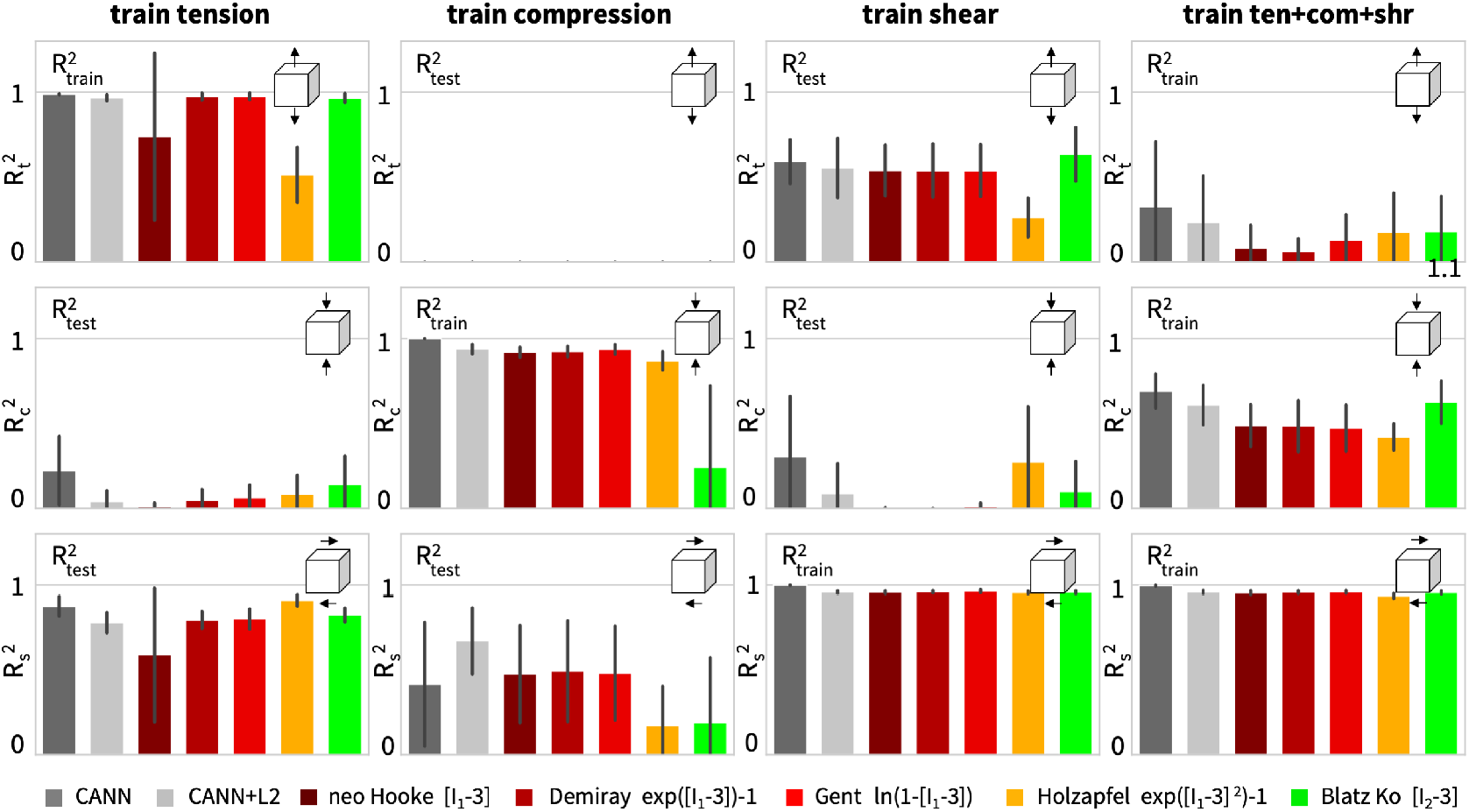
Goodness of fit for all models. Coefficients of determination *R*^2^ of our Constitutive Artificial Neural Network, without and with L2 regularization, and its special cases, the neo Hooke, Blatz Ko, Demiray, Gent, and Holzapfel models, trained with the tension, compression, and shear data [11] from Tables 1 and 2; color-coded blocks and error bars highlight the means and standard deviations of the goodness of fit *R*^2^ from 50 training runs with varying random weight initializations for each model, with colors and terms according to Figure 3.

## 4 Discussion

### Characterizing human brain tissue is a challenging but important task

Throughout the past decade, driven by the need to improve diagnostic and predictive clinical tools, neuroscience has seen an enormous, growing interest in accurately characterizing and modeling the human brain [25]. Numerous research groups have proposed competing constitutive models to best characterize the behavior of gray and white matter tissue and calibrate the model parameters in response to mechanical loading [13]. Amongst the wide variety of possible models, the neo Hooke [60], Blatz Ko [8], Mooney Rivlin [44, 52], Demiray [17], Gent [22], and Holzapfel [27] models have emerged as the most successful candidates to approximate the stress-stretch relations in the human brain. The gold standard strategy of all these approaches is to *first* select a constitutive model, either from the above list or beyond, and *then* tune its parameters by fitting the model to data. Often, these data are collected for a single loading mode–tension [43], compression [51], or shear [19]–and the parameters that fit one type of loading fail to predict the behavior for the other modes [11,45]. This simplification can have fatal consequences; for example, it could overestimate the stiffness of the brain in injury simulations. To address these limitations, our group has recently performed a comprehensive set of human brain tissue experiments in tension, compression, and shear and calibrated the neo Hooke, Mooney Rivlin, Demiray, and Gent models for four different brain regions, the cortex, basal ganglia, corona radiata, and corpus callosum [11–13]. While this approach is valuable to generate the best sets of parameters for existing models, some natural follow-up questions to ask are: How good are these models in the first place? Which one of them performs best? Are there other models that perform equally well, or even better? And, if so, how can we find them?

### Constitutive Artificial Neural Networks are a family of neural networks that a priori satisfy thermodynamic constraints

When searching for generic models that could outperform traditional constitutive models, neural networks are a natural first choice [30]. Neural networks have advanced as a powerful strategy to approximate data by cleverly combining nested and weighted activation functions with several thousand unknowns [35]. They have become the go-to strategy to interpolate data within a well-defined domain when the underlying physics are completely unknown [1]. At the same time, classical neural networks typically fail to predict the behavior outside the training domain, they violate common physical constraints, and their parameters have no real physical interpretation [37]. This has sparked the recent trend to integrate physical information into classical neural networks [36]. In the spirit of this idea, we propose a new family of neural networks that a priori satisfy common kinematic, thermodynamic, and physical constraints. Towards this goal we consult the non-linear field theories of mechanics [2, 58, 59] and constrain the network *output* to enforce thermodynamic consistency; the network *input* to enforce material objectivity, and, if desired, material symmetry and incompressibility; the *activation functions* to implement physically reasonable constitutive restrictions; and the network *architecture* to ensure polyconvexity. These ideas are not entirely new. Several recent network models are designed around enforcing thermodynamic constraints [3,24], for example through additional terms in their loss function [15]. However, the problem of overfitting sparse data with a large set of physically meaningless parameters remains [34]. This raises the questions: How do we harness decades of knowledge in constitutive modeling to create a neural network,from easy-to-understand modular building blocks, with well-defined physical parameters, that we can constrain with our domain knowledge?

### Constitutive Artificial Neural Networks can be made up of building blocks that feature prominent constitutive models

At a closer look, most popular constitutive models for human brain tissue have a similar functional structure. Here we propose to hardwire this structure into our neural network architecture. The underlying design paradigm is to *reverse-engineer* a Constitutive Artificial Neural Network that is, by design, a generalization of widely used and commonly accepted constitutive models including the neo Hooke, Blatz Ko, Mooney Rivlin, Demiray, Gent, and Holzapfel models. In Figure 3, we prototype this idea for an isotropic perfectly incompressible feed forward network with two hidden layers and four and twelve nodes. This network takes the scalar-valued first and second invariants of the deformation gradient, [*I*_1_ − 3] and [*I*_2_ − 3], as input and approximates the scalar-valued free energy function, *ψ*(*I*_1_, *I*_2_), as output. The first layer generates the first and second powers, (∘)^1^ and (∘)^2^, of the input, and the second layer applies the identity (∘), the exponential, (exp((∘)) −1), and the natural logarithm (−ln(1 − (∘))) to these powers. This results in twelve building blocks that additively feed into the final free energy function *ψ* from which we derive the Piola stress, ***P*** = *∂ψ*/*∂****F***, following standard arguments of thermodynamics. It is easy to show that our network is a *generalization* of popular constitutive models with the neo Hooke [60], Blatz Ko [8], Mooney Rivlin [44, 52], Demiray [17], Gent [22], and Holzapfel [27] models as special cases. More importantly, through a direct comparison with these models in equations (16) to (21), the weights of our network gain a clear physical interpretation. Table 7 and Figure 8 show, for example, that we recover the classical neo Hooke model with shear moduli of *μ* =1.82kPa, 0.88kPa, 0.94kPa, 0.54kPa for simultaneous training with the tension, compression, and shear data of the cortex, basal ganglia, corona radiata, and corpus callosum, which agree well with the reported values of *μ* =2.07kPa, 0.99kPa, 1.15kPa, 0.65kPa [11]. Interestingly, both our network and the parameter fit in the literature find that one of the two shear moduli of the Mooney Rivlin model is consistently zero in all four regions, while the other is *μ*_2_ = 1.88kPa, 0.90kPa, 0.97kPa, 0.56kPa for our approach compared to *μ*_1_ =2.08kPa, 1.00kPa, 1.16kPa, 0.65kPa in the literature [11]. This agrees well with other studies in which one of the Mooney Rivlin shear moduli was also significantly smaller than the other across all brain regions [45]. Figure 8 reveals several additional universal trends for human brain tissue: First, tension is not only the most challenging test to perform [43], but also the most difficult test to fit, with *R*^2^ values ranging from 0.14 to 0.48, followed by compression with 0.53 to 0.76, and shear with 0.94 to 0.96. Second, when trained simultaneously for tension, compression, and shear, all models consistently overestimate the tensile stiffness and underestimate the compression stiffness, highlighting the tension-compression asymmetry of all four types of human brain tissue [45]. Third, of all existing models, only the Holzapfel model captures the nonlinear stress response [27], suggesting that the classical invariant-based models struggle to reproduce the nonlinear behavior of human brain tissue for small deformations with 0.9 ≤ *λ* ≤ 1.1 and 0.0 ≤ *γ* ≤ 0.2. This raises the question: Can we design a Constitutive Artificial Neural Network that not only learns the best set of parameters for a given constitutive model, but also learns the model itself?

### Constitutive Artificial Neural Networks simultaneously discover both model and parameters

In essence, we propose a radically different approach towards soft tissue modeling and abandon the common strategy to *first* select a constitutive model and *then* tune its parameters by fitting the model to data [39]. Instead, we propose a family of Constitutive Artificial Neural Networks, with the general architecture in Figure 1, specified for soft tissues in Figure 3, to *simultaneously discover both, model and parameters* that best describe the data. Probing our network with the tension, compression, and shear experiments from the gray matter cortex in Table 3 and Figure 4 and from the white matter corona radiata in Table 4 and Figure 5, reveals several interesting trends: When trained with all three experiments *individually*, the network activates all its twelve terms, and fails to discover a single best model. Nonetheless, with these twelve terms, it succeeds in *interpolating* or *fitting* the training data from one experiment; however, it only performs moderately at *extrapolating* or *predicting* the test data from the other two experiments. This suggests that the data from a single loading mode are not sufficient to characterize the entire breadth of the mechanical response of human brain which agrees well with observations in the literature [11, 45]. Notably, when trained with all three experiments *simultaneously*, the Constitutive Artificial Neural Network robustly discovers a single model that best approximates the data: For the cortex, in the last columns of Table 3 and Figure 4, the network discovers four relevant terms, while the weights of the other eight terms train to zero,

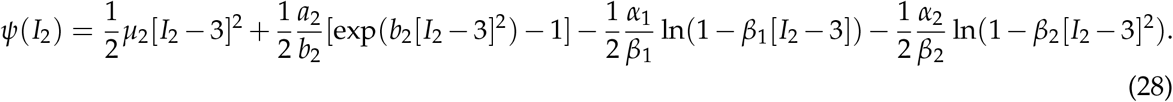

The non-zero weights translate into physically meaningful cortex parameters with well-defined physical units, the four stiffness-like parameters, *μ*_2_ = 7.60kPa, *a*_2_ = 6.23kPa, *α*_1_ = 1.25kPa, *α*_2_ = 4.67kPa, and the three nonlinearity parameters, *b*_2_ = 1.65, *β*_1_ = 0.99, *β*_2_ = 1.40. For the corona radiata, in the last columns of Table 4 and Figure 5, the network discovers four relevant terms, while the weights of the other eight terms train to zero,

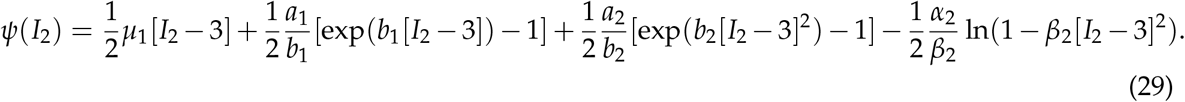

The non-zero weights translate into physically meaningful parameters with well-defined physical units, the four stiffness-like parameters, *μ*_1_ = 0.44kPa, *a*_1_ = 0.24kPa, *a*_2_ = 6.37kPa, *α*_2_ = 4.51kPa, and the three nonlinearity parameters, *b*_1_ = 0.24, *b*_2_ = 1.89, *β*_2_ = 1.18. Notably, of all seven models in Figure 9, the Constitutive Artificial Neural Network performs best in eight of all twelve cases, second best in two, and fifths in one, suggesting that it successfully discovers the model and parameters that best describe the data. Since the network *autonomously self-selects both model and parameters*, the human user no longer needs to decide which model to choose. This could have enormous implications, for example in finite element simulations: Instead of selecting a specific material model from a library of available models, finite element solvers could be designed around a single generalized model, the Constitutive Artificial Neural Network, which would *autonomously discover the model* from experimental data, populate the model parameters, and activate the relevant terms. This brings up the final and probably most interesting question: Can we learn anything from the discovery process itself?

### For human brain tissue, the Constitutive Artificial Neural Network robustly discovers *I*_2_ based models

Our Constitutive Artificial Neural Network combines the advantages of both, our knowledge of constitutive modeling [2, 6, 28, 46–48, 58] and the efficiency of neural network algorithms [33, 35, 50]. For insufficient training data that only probe individual modes, in the three left columns of Figures 4 and 5, our network approximates the overall function *ψ*(*I*_1_, *I*_2_) robustly with *R*^2^ values well above of 0.99, but similar to classical neural networks, the contributions of the individual activation functions are non-unique. Enriching the training data by multi-mode training for tension, compression, and shear in Table 5 and Figure 6 eliminates this limitation. For sufficiently rich data that probe all three modes combined, in the right columns of Figures 4 and 5, our network successfully captures the behavior of both gray and white matter, and consistently identifies the same unique subset of activation functions, *without overfitting* the data. The reduced color spectra in Figure 6 confirm that the network self-selects only a subset of activation functions, while most of its weights train to zero. For classical neural networks, a common approach to prevent overfitting is to enrich the loss function by L1 or L2 regularization as we suggest in equation (23). For L1 regularization, the discovered model and parameters are virtually identical to the plain model in Table 5 and Figure 6. For L2 regularization, the network robustly discovers a reduced model with only two terms, a subset of the non-regularized models in equations (28) and (29), while the weights of the other terms train to zero,

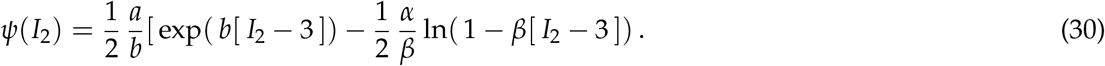

Table 6 and Figure 7 summarize the model and parameters for the regularized network with two stiffness-like parameters, *a* and *α*, and two nonlinearity parameters *b* and *β*. Strikingly, in multimode training, both the standard and L2 regularized Constitutive Artificial Neural Network consistently discover models in terms of *the second invariant only*, while all terms of the first invariant train to zero. We can easily see this selective activation in the color-coded stress terms in Figures 6 and 7, which only cold blue-type colors associated with the second invariant [*I*_2_ − 3]. The dominance of the second invariant is consistent with observations in the literature [45]. We hypothesize that the second invariant term [*I*_2_ − 3], that ranges from 0.0264 in tension to 0.0346 in compression is better suited to model the characteristic *tension-compression asymmetry* of human brain tissue than the first invariant term [*I*_1_ − 3], that only ranges only from 0.0282 to 0.0322 for our experimental range. In addition to discovering *the best model and parameters*, the goodness of fit in Figure 9 also teaches us something about *the best experiment*. If we had to select a single one experiment, tension, compression, or shear, Figure 9 suggests that the tension experiment, with the largest *R*^2^ values overall, would provide the richest data and the best insight into the behavior of human brain tissue.

### Current limitations and future applications

In the present work, we demonstrate the use of Constitutive Artificial Neural Networks for human brain under the assumption of perfect incompressibility and isotropy. The general concept extends naturally to *compressibily* or *near incompressibility* and to materials with other symmetry classes, *transverse isotropy* or *orthotropy*, by expanding the network input to other sets of strain invariants. A more dedicate extension would be to incorporate viscous effects [12], or rather *history-dependence* or *inelasticity* in general, for example, by replacing the feed forward architecture through a *long short-term memory network* with feedback connections [7], while still keeping the same overall network input, output, activation functions, and selectively connected architecture. Another limitation, which involves more complex changes, is the additive architecture of our network, which facilitates incorporating polyconvexity. Especially for human brain tissues that display a pronounced Poynting effect with shear softening in tension and shear stiffening in compression [5], it could be beneficial to introduce a *multiplicative coupling* between the individual invariants. Expressing the free energy as a truncated infinite series of products of powers of the invariants, instead of a sum of individual invariant terms, would result in a *fully connected feed forward* network architecture for which polyconvexity is cumbersome to include a priori [26]. Another technical limitation we foresee for these more complex networks, is that the majority of weights might no longer train to zero and that a more involved *L1* or *L2 regularization* could become necessary. This could artificially bias the training towards a subset of physical parameters. One interesting future direction along these lines, especially in view of human brain tissue, would be to compare *invariant-based* and *principal-stretch-based* Constitutive Artificial Neural Networks [47]. Several recent studies suggest that principal-stretch-based models outperform invariant-based models, especially in the context of combined loading and strainstifening [11,41,45]. Finally, an important extension would be to embed the network in a Bayesian inference to supplement the analysis with *uncertainty quantification* [37]. Instead of simple point estimates for the network parameters, a Bayesian Constitutive Artificial Neural Network would learn parameter distributions with means and credible intervals. In contrast to classical Bayesian Neural Networks, here, these distributions would have a clear physical interpretation, since our network weights have a well-defined physical meaning.

## 5 Conclusion

Human brain is an ultrasoft material that is difficult to test and challenging to model. Numerous competing constitutive models for human brain tissue exist in the literature, but selecting the most appropriate model remains a matter of user experience and personal preference. The underlying idea of this manuscript is to automate the process of model selection. Towards this goal, we formulate the problem of autonomous model discovery as a neural network and harness the power of gradient-based adaptive optimizers for deep learning to train the network on human brain data. However, rather than using conventional fully-connected feed-forward networks, we reverse engineer a family of Constitutive Artificial Neural Networks with a sparsely-connected architecture from a set of modular building blocks. We rationalize these building blocks from commonly accepted and widely used constitutive models for soft biological tissues, including the neo Hooke, Mooney Rivlin, Demiray, Gent, and Holzapfel models. This strategy guarantees thermodynamic consistency, material objectivity, material symmetry, physical restrictions, and polyconvexity by design. Probably even more importantly, the weights of our Constitutive Artificial Neural Networks gain a clear physical interpretation and translate naturally into common mechanical parameters. When trained with tension, compression, and shear experiments of gray and white matter tissue, the network simultaneously discovers both model and parameters, and describes the data better than any of the commonly used invariant-based models. When constrained to its individual building blocks, the network learns weights that translate into shear moduli of 1.82kPa, 0.88kPa, 0.94kPa, and 0.54kPa for the human cortex, basal ganglia, corona radiata, and corpus callosum which agree well with the reported shear moduli in these four brain regions. Taken together, Constitutive Artificial Neural Networks have the potential to enable automated model discovery and could induce a paradigm shift in soft tissue modeling, from user-defined to automated model selection and parameterization.

## Data availability

Our source code, data, and examples will be available at https://github.com/LivingMatter Lab/CANN.

## Acknowledgments

This work was supported by a DAAD Fellowship to Kevin Linka, by a National Science Foundation Graduate Research Fellowship to Sarah St. Pierre, by the Stanford School of Engineering Covid-19 Research and Assistance Fund, and by Stanford Bio-X IIP seed grant to Ellen Kuhl.

## References

[1] Alber M, Buganza Tepole A, Cannon W, De S, Dura-Bernal S, Garikipati K, Karniadakis GE, Lytton WW, Perdikaris P, Petzold L, Kuhl E (2019) Integrating machine learning and multiscale modeling: Perspectives, challenges, and opportunities in the biological, biomedical, and behavioral sciences. npj Digital Medicine 2:115.

[2] Antman SS (2005) Nonlinear Problems of Elasticity. Second edition. Springer-Verlag New York.

[3] As’ad F, Avery P, Farhat C (2022) A mechanics-informed artificial neural network approach in data-driven constitutive modeling. International Journal for Numerical Methods in Engineering 123: 2738–2759.

[4] Atanasova N, Todorovski L, Dzeroski S, Kompare B (2008) Application of automated model discovery from data and expert knowledge to a real-world domain: Lake Glumso. Ecological Modeling 212: 92–98.

[5] Balbi V, Trotta A, Destrade M, Annaidh AN (2019) Poynting effect of brain matter in torsion. Soft Matter 15:5147.

[6] Ball JM (1977) Convexity conditions and existence theorems in nonlinear elasticity. Archive for Rational Mechanics and Analysis 63: 337–403.

[7] Bhouri MA, Sahli Costabal F, Wang H, Linka K, Peirlinck M, Kuhl E, Perdikaris P (2021) COVID-19 dynamics across the US: A deep learning study of human mobility and social behavior. Computer Methods in Applied Mechanics and Engineering 382: 113891.

[8] Blatz PJ, Ko WL (1962) Application of finite elastic theory to the deformation of rubbery materials. Transactions of the Society of Rheology 6:223–251.

[9] Bongard J, Lipson H (2007) Automated reverse engineering of nonlinear dynamical systems. Proceedings of the National Academy of Sciences 104: 9943–9948.

[10] Budday S, Nay R, de Rooij R, Steinmann P, Wyrobek T, Ovaert TC, Kuhl E (2015) Mechanical properties of gray and white matter brain tissue by indentation. Journal of the Mechanical Behavior of Biomedical Materials 46: 318–330.

[11] Budday S, Sommer G, Birkl C, Langkammer C, Jaybaeck J, Kohnert Bauer M, Paulsen F, Steinmann P, Kuhl E, Holzapfel GA (2017) Mechanical characterization of human brain tissue. Acta Biomaterialia 48: 319–340.

[12] Budday S, Sommer G, Hayback J, Steinmann P, Holzapfel GA, Kuhl E (2017) Rheological characterization of human brain tissue. Acta Biomaterialia 60:315–329.

[13] Budday S, Ovaert TC, Holzapfel GA, Steinmann P, Kuhl E (2020) Fifty shades of brain: A review on the material testing and modeling of brain tissue. Archives of Computational Methods in Engineering 27:1187–1230.

[14] Center for Neurological Studies (2019) Facts about Brain Injury. https://www.neurologicstudies.com/facts-about-brain-injury.

[15] Daw A, Karpatne A, Watkins W, Read J, Kumar V (2017) Physics-guided neural networks (PGNN): An application to lake temperature modeling. arXiv. doi: 10.48550/arXiv.1710.11431.

[16] Delfino A, Stergiopulos N, Moore JE, Meister JJ (1997) Residual strain effects on the stress field in a thick wall finite element model of the human carotid bifurcation. Journal of Biomechanics 30: 777–786.

[17] Demiray H (1972) A note on the elasticity of soft biological tissues. Journal of Biomechanics 5: 309–311.

[18] Dewan MC, Rattani A, Gupta S, Baticulon RE, Hung, YC, Punchak M, Agarwal, A, Adeley AO, Shrime MG, Rubiano AM, Rosenfeld JV, Park KB (2018) Estimating the global incidence of traumatic brain injury. Journal of Neurosurgery 130(4): 1080–1097.

[19] Donnelly BR, Medige J (1997) Shear properties of human brain tissue. Journal of Biomechanical Engineering 119: 423–432.

[20] Flaschel M, Kumar S, De Lorenzis L (2021) Unsupervised discovery of interpretable hyperelastic constitutive laws. Computer Methods in Applied Mechanics and Engineering 381: 113852.

[21] GBD 2016 Traumatic Brain Injury and Spinal Cord Injury Collaborators (2019) Global, regional, and national burden of traumatic brain injury and spinal cord injury, 1990-2016: A systematic analysis for the Global Burden of Disease Study 2016. Lancet Neurology 18(1):56–87.

[22] Gent A (1996) A new constitutive relation for rubber. Rubber Chemistry and Technology 69: 59–61.

[23] Ghaboussi J, Garrett JH, Wu X (1991) Knowledge-based modeling of material behavior with neural networks. Journal of Engineering Mechanics 117: 132–153.

[24] Ghaderi A, Morovati V, Dargazany R (2020) A physics-informed assembly for feed-forward neural network engines to predict inelasticity in cross-linked polymers. Polymers 12: 2628.

[25] Goriely A, Geers MGD, Holzapfel GA, Jayamohan J, Jerusalem A, Sivaloganathan S, Squier W, van Dommelen JAW, Waters S, Kuhl E (2015) Mechanics of the brain: Perspectives, challenges, and opportunities. Biomechanics Modeling and Mechanobiology. 14: 931–965.

[26] Hartmann S, Neff P (2003) Polyconvexity of generalized polynomial-type hyperelastic strain energy functions for near-incompressibility. International Journal of Solids and Structures 40: 2767–2791.

[27] Holzapfel GA, Gasser TC, Ogden RW (2000) A new constitutive framework for arterial wall mechanics and comparative study of material models. Journal of Elasticity 61:1–48.

[28] Holzapfel GA (2000) Nonlinear Solid Mechanics: A Continuum Approach to Engineering. John Wiley & Sons, Chichester.

[29] Holzapfel GA, Linka K, Sherifova S, Cyron C (2021) Predictive constitutive modelling of arteries by deep learning. Journal of the Royal Socienty Interface 18:20210411.

[30] Hopfield JJ (1982) Neural networks and physical systems with emergent collective computational abilities. Proceedings of the National Academy of Science 79:2554–2558.

[31] Hoppstadter M, Pullmann D, Seydewitz R, Kuhl E, Bol M (2022) Correlating the microstructural architecture and macrostructural behaviour of the brain. Acta Biomaterialia 151:379–395.

[32] Kakaletsis S, Lejeune E, Rausch MK (2022) Can machine learning accelerate soft material parameter identification from complex mechanical test data? Biomechanics and Modeling in Mechanobiology. doi:10.1007/s10237-022-01631-z.

[33] Karniadakis GE, Kevrekidis IG, Lu L, Perdikaris P, Wang S, Yang L (2021) Physics-informed machine learning. Nature Reviews Physics 3:422–440.

[34] Klein DK, Fernandez M, Martin RJ, Neff P, Weeger O (2022) Polyconvex anisotropic hyperelasticity with neural networks. Journal of the Mechanics and Physics of Solics 159: 105703.

[35] LeCun Y, Bengio Y, Hinton G (2015) Deep learning. Nature 521: 436–444.

[36] Linka K, Hillgartner M, Abdolazizi KP, Aydin RC, Itskov M, Cyron CJ (2021) Constitutive artificial neural networks: A fast and general approach to predictive data-driven constitutive modeling by deep learning. Journal of Computational Physics 429: 110010.

[37] Linka K, Schafer A, Meng X, Zou Z, Karniadakis GE, Kuhl E (2022) Bayesian Physics-Informed Neural Networks for real-world nonlinear dynamical systems. Computer Methods in Applied Mechanics and Engineering; online first doi:10.1016/j.cma.2022.115346.

[38] Linka K, Cavinato C, Humphrey JD, Cyron CJ (2022) Predicting and understanding arterial elasticity from key microstructural features by bidirectional deep learning by deep learning. Acta Biomaterialia 147: 63–72.

[39] Linka K, Kuhl E (2022) A new family of Constitutive Artificial Neural Networks towards automated model discovery. Computer Methods in Applied Mechanics and Engineering. doi: 10.1016/j.cma.2022.115731.

[40] Masi F, Stefanou I, Vannucci P, Maffi-Berthier V (2021) Thermodynamics-based artificial neural networks for constitutive modeling. Journal of the Mechanics and Physics of Solids 147: 04277.

[41] Mihai LA, Chin L, Janmey PA, Goriely A (2015) A comparison of hyperelastic constitutive models applicable to brain and fat tissues. Journal of the Royal Society Interface 12: 20150486.

[42] Mihai LA, Budday S, Holzapfel GA, Kuhl E, Goriely A (2017) A family of hyperelastic models for human brain tissue. Journal of the Mechanics and Physics of Solids 106: 60–79.

[43] Miller K, Chinzei K (2002) Mechanical properties of brain tissue in tension. Journal of Biomechanics 35: 483–490.

[44] Mooney M (1940) A theory of large elastic deformations. Journal of Applied Physics 11: 582–590.

[45] Moran R, Smith JH, Garcia JJ (2014) Fitted hyperelastic parameters for human brain tissue from reported tension, compression, and shear tests. Journal of Biomechanics 47: 3762–3766.

[46] Noll W (1958) A mathematical theory of the mechanical behavior of continuous media. Archive of Rational Mechanics Analysis 2: 197–226.

[47] Ogen RW (1972) Large deformation isotropic elasticity – on the correlation of theory and experiment for incompressible rubberlike solids. Proceedings of the Royal Socienty London Series A 326: 565–584.

[48] Planck M (1897) Vorlesungen über Thermodynamik. Verlag von Veit & Comp, Leipzig.

[49] Prevost TP, Balakrishnan A, Suresh S, Socrate S (2011) Biomechanics of brain tissue. Acta Biomaterialia 7:83–95.

[50] Raissi M, Perdikaris P, Karniadakis GE (2019) Physics-informed neural networks: a deep learning framework for solving forward and inverse problems involving nonlinear partial differential equations. Journal of Computational Physics 378:686–707.

[51] Rashid B, Destrade M, Gilchrist MD (2012) Mechanical characterization of brain tissue in compression at dynamic strain rates. Journal of the Mechanical Behavior of Biomedical Materials 10: 23–38.

[52] Rivlin RS (1948) Large elastic deformations of isotropic materials. IV. Further developments of the general theory. Philosophical Transactions of the Royal Society of London Series A 241: 379–397.

[53] Rivlin RS, Saunders DW (1951) Large elastic deformations of isotropic materials. VII. Experiments on the deformation of rubber. Philosophical Transactions of the Royal Society of London Series A 243: 251–288.

[54] Schmidt M, Lipson H (2009) Distilling free-form natural laws from experimental data. Science 324: 81–85.

[55] Schulte R, Karca C, Ostwald R, Menzel A (2022) Machine learning-assisted parameter identification for constitutive models based on concatenated normalised modes. European Journal of Mechanics A/Solids.

[56] Shen Y, Chandrashekhara K, Breig WF, Oliver LR (2004) Neural Network based constitutive model for rubber material. Rubber Chemistry and Technology 77: 257–277.

[57] Tac V, Sahli Costabal F, Buganza Tepole A. Data-driven tissue mechanics with polyconvex neural ordinary differential equations. Computer Methods in Applied Mechanics and Engineering 398: 115248.

[58] Truesdell C, Noll W (1965) Non-linear field theories of mechanics. In: Flügge S, Ed., Encyclopedia of Physics, Vol. III/3, Spinger, Berlin.

[59] Truesdell C (1969) Rational Thermodynamics, Lecture 5. McGraw-Hill, New York.

[60] Treloar LRG (1948) Stresses and birefringence in rubber subjected to general homogeneous strain. Proceedings of the Physical Society 60: 135–144.

[61] Weickenmeier J, de Rooij R, Budday S, Steinmann P, Ovaert TC, Kuhl E (2016) Brain stiffness increases with myelin content. Acta Biomaterialia 42:265–272.

[62] Weickenmeier J, Kurt M, Ozkaya E, Wintermark M, Butts Pauly K, Kuhl E (2018) Magnetic resonance elastography of the brain: A comparison between pigs and humans. Journal of the Mechanical Behavior of Biomedical Materials 77:702–710.

[63] Zopf C, Kaliske M (2017) Numerical characterisation of uncured elastomers by a neural network based approach. Computers and Structures 182: 504–525.

